# Comparative Metabolomics and Lipidomics Reveal Selective Fatty Acyl Partitioning and Metabolic Reorganization Associated with Long-Term Cold Adaptation in *Euphausia superba*

**DOI:** 10.64898/2026.11.10.765018

**Authors:** Syuan-Huai Shih, Jean-Yves Toullec, Chi-Ying Lee

## Abstract

Antarctic krill (*Euphausia superba*) inhabits the persistently cold waters of the Southern Ocean, where lipid metabolism supports membrane function and energy storage. Despite extensive characterization of euphausiid lipid classes and bulk fatty acids, fatty acyl organization across lipid pools remains poorly understood. We integrated untargeted metabolomics and lipidomics to compare whole-body biochemical profiles of *E. superba* and its congener *E. triacantha*. Metabolomics revealed differences in glycerophospholipid (GP) and glycerolipid (GL) metabolism, together with alterations in pathways associated with nucleotide metabolism, redox regulation, and lipid precursor supply. Lipidomics further revealed selective remodeling at the lipid-class and molecular-species levels rather than a global increase in lipid unsaturation. Relative to *E. triacantha*, *E. superba* showed lower relative abundances of multiple lysophospholipid classes and a higher relative abundance of cardiolipin. *E. superba* also showed higher relative abundances of polyunsaturated glycerophospholipids, whereas a subset of predominantly polyunsaturated triacylglycerol species showed lower relative abundances. C20:5, C22:5, and C22:6 showed higher relative abundances in the GP pool but lower relative abundances in the GL pool of *E. superba*. Together, these findings place selective fatty acyl partitioning within an integrated metabolic framework associated with long-term cold adaptation in *E. superba*.

## Introduction

Antarctic krill (*Euphausia superba*) is a central component of the Southern Ocean ecosystem, sustaining one of the world’s most productive marine food webs. With an estimated circumpolar biomass of approximately 379 million metric tons (Atkinson et al., 2009), *E. superba* serves as a major trophic link connecting microalgae to a wide range of upper-trophic predators, including baleen whales, seals, and penguins (Murphy et al., 2007). Its substantial lipid content, which can exceed 30% of dry body mass (Clarke, 1984; Hagen et al., 2001), contributes to its value as an energy-rich resource for higher-trophic-level predators. Antarctic krill also plays an important role in Southern Ocean biogeochemistry through diel vertical migration, production of rapidly sinking fecal pellets, and nutrient recycling, thereby supporting regional carbon export and the biological carbon pump (Cavan et al., 2019; Pauli et al., 2021; Smith et al., 2025; Tovar-Sanchez et al., 2007).

Despite its ecological importance, *E. superba* faces increasing pressure from rapid ocean warming, declining sea-ice cover, and expanding commercial exploitation (Flores et al., 2012; Kawaguchi et al., 2024; Savoca et al., 2024). As a stenothermal species adapted to persistently cold Antarctic conditions, *E. superba* must sustain cellular function and metabolic homeostasis at low temperature (Hagen et al., 2001; Meyer, 2012). Understanding the physiological basis of this cold specialization is therefore important not only for explaining the biochemical basis of Antarctic adaptation, but also for evaluating how this species may respond to ongoing environmental change.

Lipid metabolism is likely to play a central role in this physiological specialization because lipids must simultaneously support membrane function and energy storage under persistent low-temperature conditions (Hazel and Williams, 1990; Wu et al., 2023). *E. superba* contains abundant glycerophospholipids (GPs), particularly phosphatidylcholine, together with substantial triacylglycerol (TG) stores and high levels of polyunsaturated fatty acids such as eicosapentaenoic acid and docosahexaenoic acid (Auerswald et al., 2009; Saether et al., 1985; Hagen et al., 2001). Previous studies have documented substantial variation in euphausiid lipid composition across species, habitats, nutritional states, seasons, and life-history stages, including pronounced differences in the relative contributions of TGs, wax esters, and phospholipids (Bottino, 1975; Hagen et al., 2001; Hagen and Kattner, 1998; Hellessey et al., 2018; Ju et al., 2009; Mayzaud et al., 2003; Saether et al., 1986). Most euphausiid lipid studies have nevertheless focused on total lipid abundance, lipid-class composition, or bulk fatty acid profiles (Hagen et al., 2001; Ju et al., 2009; Mayzaud et al., 2003; Phleger et al., 2002; Saether et al., 1986; Murzina et al., 2023), although recent work has begun to resolve intact phospholipid molecular composition in specific tissues of *E. superba* (Heyen et al., 2023). Because the metabolic and structural roles of fatty acids depend partly on the lipid pools into which they are incorporated (Saether et al., 1986), bulk measurements may obscure functionally important differences in molecular lipid organization. Consequently, how specific fatty acyl chains are organized across GP and glycerolipid (GL) pools, and how this molecular organization is integrated with metabolic pathways during long-term cold adaptation, remain poorly understood.

Comparative analyses of closely related species occupying contrasting thermal environments provide a useful framework for addressing these questions. The congeneric species *E. superba* and *E. triacantha* offer such a comparison: *E. superba* is distributed predominantly in Antarctic waters, whereas *E. triacantha* has a more northerly distribution spanning Antarctic and sub-Antarctic waters (Cuzin-Roudy et al., 2014). This contrast allows us to ask whether adaptation to distinct thermal niches is associated with differences in lipid abundance, molecular lipid organization, and metabolic profiles.

Here, we combined untargeted metabolomics and lipidomics to characterize whole-body biochemical differences between *E. superba* and *E. triacantha*. Metabolomic analyses characterized pathway-level metabolic differences, whereas lipidomic analyses examined GP and GL pools at the levels of molecular composition, lipid unsaturation, and fatty acyl chain distribution. Our results reveal selective partitioning of polyunsaturated fatty acyl chains across GP and GL pools in *E. superba*, accompanied by metabolic reorganization. Together, these findings place molecular lipid remodeling within an integrated metabolic framework associated with long-term cold adaptation in Antarctic krill.

## Materials and Methods

### Animal Collection

*E. superba* were fished off the French Antarctic station Dumont d’Urville (DDU) in Terre Adélie just beyond the continental plateau (65.30° S, 139.57° E, 01/2011). Fishing was carried out using an Isaacs–Kidd midwater trawl (IKMT) deployed from the research vessel *L’Astrolabe*. Sampling depth was determined by sonar observations of krill swarms, and specimens were collected from swarms occurring at depths between 20 and 50 m. *E. triacantha* were fished in the Indian Ocean during the Obs Austral/MD206 cruise aboard the research vessel *Marion Dufresne* (49°12 ′ S, 69°49 ′ E, 01/2017), using an IKMT during a trawl at a depth of 1,000 meters. (TOULLEC Jean-Yves, KOUBBI Philippe (2017) VT 155 / REPCCOAI cruise, RV Marion Dufresne, https://doi.org/10.17600/17017100). Immediately after collection, specimens of both species were frozen in liquid nitrogen and maintained under liquid-nitrogen conditions during transport to the laboratory. Samples were subsequently stored at −80 °C until metabolomic and lipidomic analyses. Six independent biological specimens from each species were used for subsequent analyses.

### Untargeted Metabolomic and Lipidomic Profiling by UHPLC–MS/MS

Whole-body metabolite extraction and mass spectrometry analysis were performed with minor modifications based on previously published protocols (Dunn et al., 2011; Want et al., 2010). The specific modifications are described below.

#### Whole-Body Metabolite Extraction

Each whole krill specimen was ground to a fine powder under liquid nitrogen using a pre-cooled mortar and pestle. 50 mg of the resulting whole-body powder was then transferred into a microcentrifuge tube containing 1,000 μL of pre-chilled (–20 °C) extraction solvent (methanol: acetonitrile: water, 2:2:1, v/v/v) supplemented with 2-chloro-L-phenylalanine (5 μg/mL) as an internal standard. Samples were homogenized with ceramic beads using a bead mill homogenizer (JXFSTPRP-24, Shanghai Jingxin Industrial Development Co., Shanghai, China) at 45 Hz for 4 min, followed by sonication (PS-60AL, Shenzhen Leidabang Electronic Co., Ltd., Shenzhen, China) for 5 min in an ice-water bath. The homogenization and sonication steps were performed for three cycles. All procedures were performed on ice unless otherwise specified. The extracts were incubated at –20 °C for 1 h to facilitate protein precipitation and centrifuged at 12000 rpm for 15 min at 4 °C using a Heraeus Fresco 17 centrifuge (Thermo Fisher Scientific, Bremen, Germany). An aliquot of supernatant (500 μL) was transferred to fresh tubes and evaporated to dryness using a vacuum concentrator (SpeedVac, Thermo Fisher Scientific) without heating. The dried residues were redissolved in 100 μL of acetonitrile:water (1:1, v/v), vortexed for 30 s, sonicated for 10 min in an ice-water bath, and centrifuged again at 12000 rpm for 15 min at 4 °C. Finally, 60 μL of the supernatant was transferred into LC–MS glass vials for UHPLC–QTOF–MS analysis. **Lipid Extraction**

For lipidomic analysis, lipids were extracted using a methyl tert-butyl ether (MTBE)-based extraction method. Briefly, 50 mg of the resulting whole-body powder was mixed with 200 μL of water and 480 μL of MTBE:methanol (5:1, v/v), followed by homogenization and sonication as described above. After incubation at –20 °C for 1 h, samples were centrifuged at 13,000 rpm for 15 min at 4 °C to facilitate phase separation. The upper organic phase (380 μL) was collected and evaporated to dryness under vacuum. The dried extracts were redissolved in 100 μL of dichloromethane:methanol (1:1, v/v), vortexed for 30 s, sonicated for 10 min in an ice-water bath, and centrifuged again at 12,000 rpm for 15 min at 4 °C. Finally, 60 μL of the supernatant was transferred into LC–MS glass vials for UHPLC–QTOF–MS lipidomic analysis.

#### Mass-Spectrometry-Based Analysis

Metabolomic profiling was performed using an ultra-high-performance liquid chromatography (UHPLC) system (1290 Infinity, Agilent Technologies, Santa Clara, CA, USA) coupled to a TripleTOF 6600 mass spectrometer (SCIEX, Framingham, MA, USA). Chromatographic separation was achieved on an ACQUITY UPLC BEH Amide column (1.7 μm, 2.1 × 100 mm; Waters, Milford, MA, USA) maintained at 25 °C. The mobile phase consisted of solvent A (25 mM ammonium acetate and 25 mM ammonium hydroxide in water, pH 9.75) and solvent B (acetonitrile). The gradient elution was programmed as follows: 0 min, 95% B; 0.5 min, 95% B; 7 min, 65% B; 8 min, 40% B; 9 min, 40% B; 9.1 min, 95% B; and 12 min, 95% B. The flow rate was 0.5 mL/min, and the injection volume was 2 μL. Mass spectrometric detection was performed in both positive and negative electrospray ionization (ESI) modes. MS/MS spectra were acquired in information-dependent acquisition (IDA) mode using Analyst TF 1.7 software (SCIEX). TOF-MS survey scans were acquired from m/z 60 to 1200, and MS/MS product-ion scans were acquired from m/z 25 to 1200. In each IDA cycle, precursor ions with intensities greater than 100 were selected for MS/MS fragmentation. Up to 15 MS/MS spectra were acquired per cycle at a collision energy of 30 eV, with an accumulation time of 50 ms for each spectrum. ESI source parameters were as follows: ion source gas 1 (GS1), 60 psi; ion source gas 2 (GS2), 60 psi; curtain gas, 35 psi; source temperature, 650 °C; ion spray voltage floating (ISVF), 5,000 V (positive mode) or –4,000 V (negative mode). A pooled quality-control (QC) sample was prepared by mixing 10 μL aliquots from each extract, and 60 μL of the pooled mixture was transferred to an LC–MS vial for analysis. Three QC injections were included in the analytical batch to assess analytical reproducibility. The reproducibility of the internal standard 2-chloro-L-phenylalanine was evaluated across all QC injections, with relative standard deviations (RSDs) of 3.38% and 2.27% in positive and negative ion modes, respectively. The measured m/z values of 2-chloro-L-phenylalanine was 200.0479 in positive ion mode and 198.0358 in negative ion mode, with retention times (RT) of 3.68 min and 3.70 min, respectively.

Lipidomic profiling was performed using the same UHPLC and mass spectrometer setup. Chromatographic separation was achieved on a Phenomenex Kinetex C18 column (1.7 μm, 100 × 2.1 mm; Phenomenex, Torrance, CA, USA) maintained at 55 °C. The mobile phase consisted of solvent A (10 mM ammonium formate in water/acetonitrile, 40:60, v/v) and solvent B (10 mM ammonium formate in acetonitrile/isopropanol, 10:90, v/v). The gradient program was as follows: 0 min, 40% B; 12 min, 100% B; 13.5 min, 100% B; 13.7 min, 40% B; and 18 min, 40% B. The flow rate was 0.3 mL/min, and the injection volume was 0.5 μL in positive ion mode and 2 μL in negative ion mode. MS/MS spectra were acquired in IDA mode using Analyst TF 1.7 software (SCIEX). TOF-MS survey scans were acquired from m/z 200 to 2000, and MS/MS product-ion scans were acquired from m/z 100 to 2000. In each IDA cycle, precursor ions with intensities greater than 100 were selected for MS/MS fragmentation. Up to 15 MS/MS spectra were acquired per cycle at a collision energy of 35 eV, with an accumulation time of 50 ms for each spectrum. Mass spectrometric detection was performed in both positive and negative ESI modes. The ESI source parameters were as follows: ion source gas 1 (GS1), 60 psi; ion source gas 2 (GS2), 60 psi; curtain gas, 30 psi; source temperature, 550 °C; ion spray voltage floating (ISVF), 5,500 V (positive mode) or –4,500 V (negative mode). Pooled QC samples were prepared as described for metabolomics.

#### Data Preprocessing and Annotation

Data processing and annotation were performed with reference to Wang et al., 2016. Raw MS data (.wifffiles) were converted to mzXML format using ProteoWizard and processed with the R package XCMS (version 3.2) (Smith et al., 2006) to generate a data matrix containing retention time (RT), mass-to-charge ratio (m/z), and peak intensity. CAMERA (Kuhl et al., 2012) was used for peak annotation. Additional filtering was performed using an in-house data-processing workflow. A minfrac value of 0.5 was applied, requiring a peak to be detected in at least 50% of samples within a group for retention, and a cutoff value of 0.6 was applied during pseudospectrum processing. Metabolite identification was performed using an in-house MS/MS spectral database generated from purchased reference standards analyzed under the same LC–MS conditions. Peak areas were integrated for annotated metabolites, and relative abundances were calculated using peak-area normalization (Qu et al., 2015).

#### Metabolome data analysis

Metabolites detected only at the MS1 level without corresponding MS/MS support were excluded from downstream analyses to increase annotation confidence. For each sample, the relative abundance of each retained metabolite was calculated as follows:

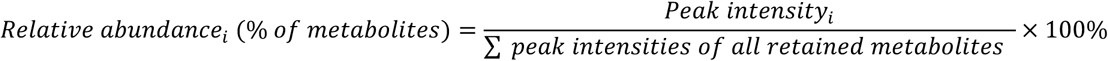

Metabolic pathway enrichment analysis was performed using MetaboAnalyst 6.0 (Pang et al., 2024). All 577 MS/MS-supported metabolites were included in the analysis and mapped to *Drosophila melanogaster* pathways in the Kyoto Encyclopedia of Genes and Genomes (KEGG) database (accessed February 2026). Prior to pathway analysis, metabolite abundance data were subjected to median normalization, log2 transformation, and Pareto scaling in MetaboAnalyst 6.0. Pathways with Holm-adjusted *p* < 0.05 were considered significantly enriched. The ten significantly enriched pathways with the lowest Holm-adjusted *p*-values were presented in Figure 1B, and all significantly enriched pathways are shown in Figure S2. Representative lipid-related pathways were additionally visualized using heatmaps of log2-transformed metabolite relative abundances. All metabolome data, including the filtered metabolite list, peak intensities, and calculated relative abundances, are provided in the Supplemental Information.

**Figure 1.**
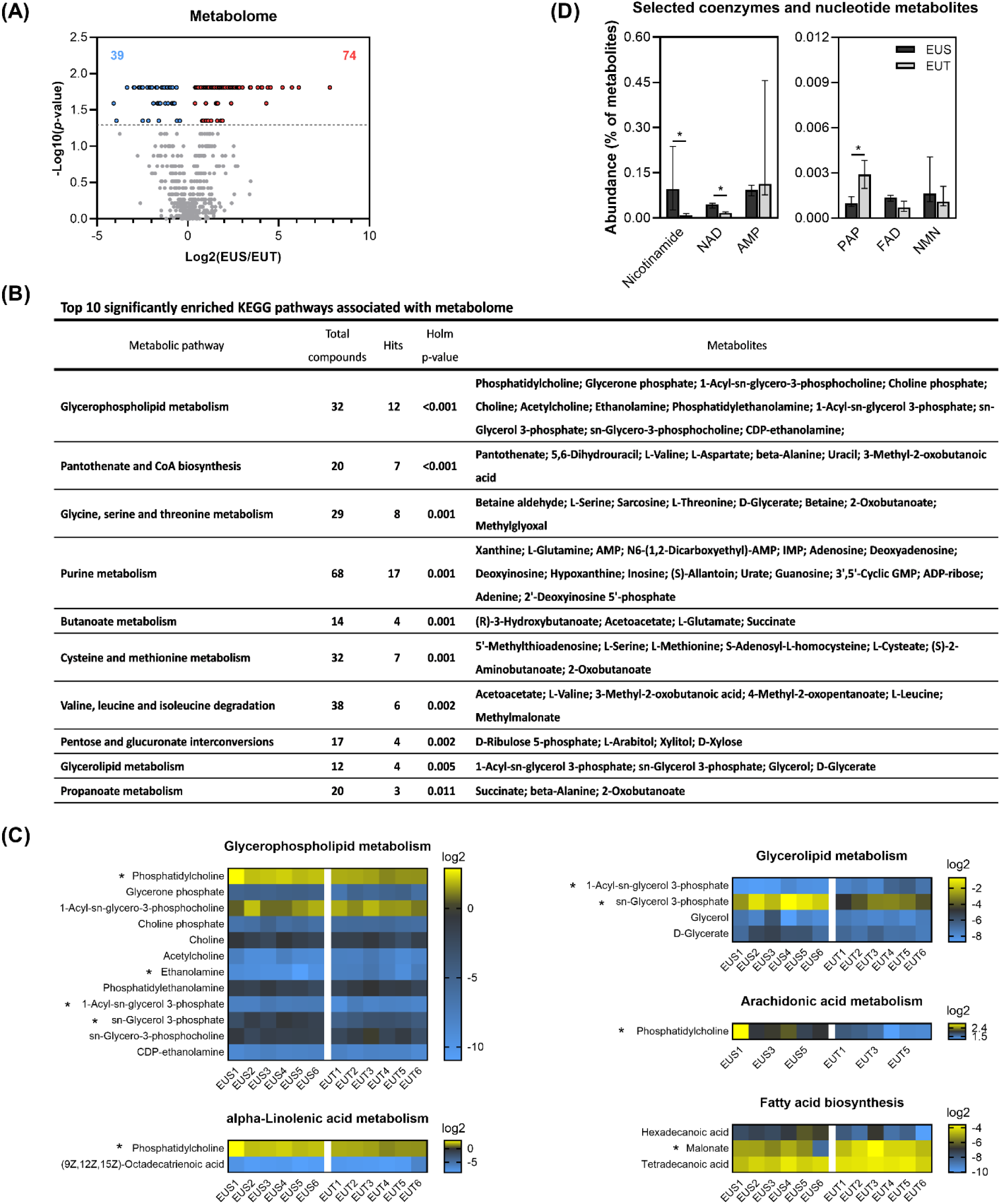
Global metabolomic differences between *E. superba* and *E. triacantha* and associated enriched metabolic pathways. (A) Differential abundance of metabolites detected by LC–MS/MS. Metabolite differences are displayed as the log2 fold change of *E. superba* (EUS) relative to *E. triacantha* (EUT) against the negative log10-transformed FDR-adjusted *p*-value based on two-sided Mann– Whitney U tests. The horizontal dotted line indicates the significance threshold (FDR < 0.05). Red and blue points indicate metabolites with significantly higher relative abundances in EUS and EUT, respectively; numbers indicate the corresponding metabolite counts. (B) The ten significantly enriched KEGG pathways with the lowest Holm-adjusted *p*-values. All significantly enriched pathways are shown in Figure S2. (C) Heatmaps showing the relative abundances of metabolites associated with representative lipid-related enriched pathways. Values represent log2-transformed metabolite relative abundances. Each column corresponds to one biological replicate of EUS or EUT. Asterisks indicate metabolites that differed significantly between *E. superba* and *E. triacantha* (FDR < 0.05). (D) Relative abundances of selected coenzyme- and nucleotide-related metabolites in EUS and EUT. Data are presented as medians with interquartile ranges (IQRs), with six biological replicates per species. Black and gray bars represent EUS and EUT, respectively. * indicates FDR < 0.05 based on two-sided Mann–Whitney U tests with Benjamini–Hochberg correction. Abbreviations: NAD, nicotinamide adenine dinucleotide; AMP, adenosine monophosphate; PAP, adenosine 3′,5′-diphosphate; FAD, flavin adenine dinucleotide; NMN, nicotinamide mononucleotide.

#### Lipidome data analysis

Lipid species were classified according to LIPID MAPS nomenclature (https://www.lipidmaps.org). Only lipid annotations supported by MS/MS spectra were retained for downstream analysis. Individual lipid molecular species were reported at the fatty acyl composition level (e.g., PC(24:4/22:5)). The relative abundance of each lipid species was calculated as follows:

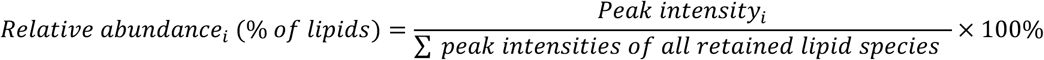

Lipid molecular species were classified into saturated (SA), monounsaturated (MU), and polyunsaturated (PU) according to the degree of unsaturation of their constituent fatty acyl chains. Species containing at least one polyunsaturated fatty acyl chain (>1 double bond) were classified as PU, whereas species containing one or more monounsaturated fatty acyl chains (1 double bond) but no polyunsaturated chains were classified as MU. Species composed exclusively of saturated fatty acyl chains (0 double bonds) were classified as SA. Accordingly, PC(18:1/18:1) was classified as MU, whereas PC(16:0/20:5) was classified as PU.

For fatty acyl composition analysis, each lipid molecular species was decomposed into its constituent fatty acyl chains. The relative abundance of a lipid species was assigned to each constituent fatty acyl chain according to its stoichiometric occurrence. Thus, when the same fatty acyl chain occurred more than once within a lipid species, its contribution was counted accordingly; for example, the contribution of 18:1 from PC(18:1/18:1) was calculated as twice the relative abundance of that lipid species. Contributions from identical fatty acyl chains across all lipid molecular species, irrespective of lipid category, were then summed, and the total fatty acyl abundance within each sample was normalized to 100%. Fatty acyl contributions were subsequently examined according to the lipid category from which they were derived.

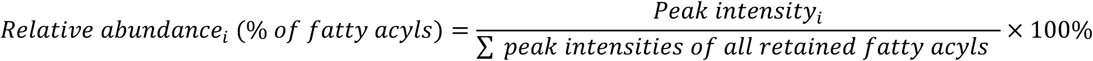

For analyses based on total carbon number and total double-bond number, the carbon atoms and double bonds of the constituent fatty acyl chains were summed within each lipid molecular species. For example, PC(24:4/22:5) was represented as PC(46:9). Lipid molecular species were subsequently grouped according to lipid class, the SA/MU/PU classification described above, or summed carbon and double-bond number, as appropriate for each analysis. Fatty acyl composition data were analyzed separately according to individual chain identity and chain length.

### Statistical Analysis

Data are presented as medians with interquartile ranges (IQRs), with six biological replicates per krill species. Differences between *E. superba* and *E. triacantha* were assessed using two-sided Mann–Whitney U tests (Mann and Whitney, 1947). For feature-level differential abundance analyses of metabolites and lipid molecular species, *p*-values were adjusted using the Benjamini–Hochberg procedure (Benjamini and Hochberg, 1995), with FDR < 0.05 considered statistically significant. Analyses of lipid-category, lipid-class, and aggregated unsaturation-category abundances were evaluated using nominal Mann– Whitney U-test *p*-values, with significance thresholds specified in the corresponding figure legends. For fatty acyl composition analyses, interspecific differences were assessed using two-sided Mann–Whitney U tests, and Benjamini–Hochberg-adjusted FDR values were additionally calculated across fatty acyl comparisons; fatty acyl chains remaining significant at FDR < 0.05 were specifically indicated in Figure 4.

Principal component analysis (PCA) was performed separately for the metabolomic and lipidomic datasets using the prcomp function in the R stats package. Prior to PCA, each dataset was independently subjected to median normalization, log2(x + 1) transformation, and Pareto scaling (mean-centering followed by division by the square root of the standard deviation). PCA was performed on the resulting preprocessed matrices without additional centering or scaling. Differences in multivariate composition between species were assessed separately for the metabolomic and lipidomic datasets by permutational multivariate analysis of variance (PERMANOVA) using adonis2 in the vegan package, based on Euclidean distances with 999 permutations. PCA plots were generated using factoextra and ggplot2 and assembled using patchwork. Homogeneity of multivariate dispersion was assessed using PERMDISP with the betadisper function in the vegan package, based on distances to group spatial medians and the same Euclidean distance matrices used for PERMANOVA. Significance was assessed using 999 permutations.

Spearman’s rank correlation analysis was performed among the 113 differential metabolites and 218 differential lipid species identified between *E. superba* and *E. triacantha*, using all 12 biological samples. Pairwise correlations were calculated between individual lipid species and metabolites and among lipid species. Correlations with |*r*| ≥ 0.9 and FDR < 0.05 were retained for network analysis. For visualization, significant lipid–metabolite correlations were aggregated by lipid class, with metabolites represented as a single group, whereas significant lipid–lipid correlations were aggregated according to the lipid classes of both correlated species. Ribbon width represents the number of retained pairwise correlations between the corresponding groups, whereas ribbon color represents the direction of the mean Spearman correlation coefficient across the aggregated pairs, with red and blue indicating positive and negative mean correlations, respectively. Chord diagrams were generated using the circlize package in R.

Data processing and statistical analyses were performed using R (version 4.5.2) and Python (version 3.14.2). Custom Python scripts were used for downstream processing of annotated lipid molecular species, including classification according to constituent fatty acyl unsaturation, fatty acyl composition analysis, and calculation of summed carbon and double-bond numbers. Microsoft Excel was used for data organization where applicable. Graphical outputs were generated using R and GraphPad Prism (GraphPad Software, La Jolla, CA, USA) and assembled for publication using Adobe Illustrator (Adobe Inc., San Jose, CA, USA).

## Results

### Metabolic Pathway Divergence between *E. superba* and *E. triacantha*

Untargeted metabolomic analysis using liquid chromatography-tandem mass spectrometry (LC-MS/MS) identified a total of 577 metabolites across the metabolomes of *E. superba* and *E. triacantha*. Principal component analysis (PCA) revealed a clear separation between the metabolomic profiles of the two krill species (PERMANOVA, R^2^ = 0.316, *p* = 0.002; **Figure S1A**), with no significant difference in multivariate dispersion detected between species (PERMDISP, *p* = 0.116). PC1 and PC2 explained 34.3% and 22.1% of the total variance, respectively. Differential abundance analysis further revealed that 113 metabolites differed significantly between the two krill species (FDR < 0.05), of which 74 showed higher abundance and 39 lower abundance in *E. superba* relative to *E. triacantha* **(Figure 1A)**.

To characterize the pathway-level differences associated with these detected metabolites, enrichment analysis was performed against the Kyoto Encyclopedia of Genes and Genomes (KEGG) database using the MetaboAnalyst platform. In total, 57 pathways were mapped, of which 17 were significantly enriched (Holm-adjusted *p* < 0.05). The top 10 enriched pathways were presented in **Figure 1B**, and the complete set of significantly enriched pathways was provided in **Figure S2**. Five of the 17 enriched pathways were related to lipid metabolism, including glycerophospholipid metabolism, glycerolipid metabolism, arachidonic acid metabolism, alpha-linolenic acid metabolism, and fatty acid biosynthesis (**Figure 1B and S2**). Several pathway-associated metabolites differed significantly between the two species (**Figure 1C**). Specifically, the abundances of phosphatidylcholine, 1-acyl-*sn*-glycerol 3-phosphate, and *sn*-glycerol 3-phosphate were significantly higher in *E. superba* (FDR < 0.05), whereas ethanolamine and malonic acid were significantly lower.

In addition to lipid-related pathway differences, alterations were also observed in coenzyme and nucleotide-related metabolism, including pantothenate and CoA biosynthesis, purine metabolism, pentose and glucuronate interconversions, and the pentose phosphate pathway **(Figure 1B and S2)**. Furthermore, selected coenzyme-and nucleotide-related metabolites showed distinct abundance patterns, with *E. superba* exhibiting significantly higher levels of nicotinamide (NAM) and nicotinamide adenine dinucleotide (NAD), along with lower levels of adenosine 3’,5’-diphosphate (PAP), compared with *E. triacantha* **(Figure 1D)**.

Together, these results reveal distinct pathway-level metabolic difference between *E. superba* and *E. triacantha*, involving lipid-, nucleotide-, and coenzyme-related metabolism.

### Class-Level Remodeling of Glycerophospholipids in *E. superba* Compared with *E. triacantha*

To further characterize the lipidomic differences between the two krill species, untargeted LC–MS/MS-based lipidomic analysis was performed on whole-body samples of *E. superba* and *E. triacantha.* Prior to lipidomic analysis, free fatty acids (FFAs) detected in the metabolomic profiles were quantified as an indicator of potential lipid degradation during sample processing (Clarke, 1980). Individual FFA species remained at low relative abundances (<1% of total metabolite abundance; **Figure S3**), providing no indication of extensive lipid degradation during sample processing.

A total of 859 lipid species were identified across the two krill lipidomes. PCA revealed a clear separation between the lipidomic profiles of *E. superba* and *E. triacantha* (PERMANOVA, R^2^ = 0.386, *p* = 0.013; **Figure S1B**), with no significant difference in multivariate dispersion detected between species (PERMDISP, p = 0.195). PC1 and PC2 explained 57.5% and 16.2% of the total variance, respectively. The identified lipids were assigned to three major lipid categories, glycerophospholipids (GP), glycerolipids (GL), and sphingolipids (SP), together with cholesteryl esters (CE). The three major lipid categories were further subdivided into 24 lipid classes. Abbreviations and the number of lipid species identified within each class were summarized in **Figures 2A and S4**. GP and GL represented the dominant lipid categories in both krill lipidomes, whereas SP and CE together accounted for less than 1% of total lipid abundance (**Figure 2B)**. Because GP and GL represented the dominant lipid categories and their corresponding metabolic pathways were significantly enriched in the metabolomic analysis (**Figure 1B)**, subsequent analyses focused primarily on these two lipid categories.

**Figure 2.**
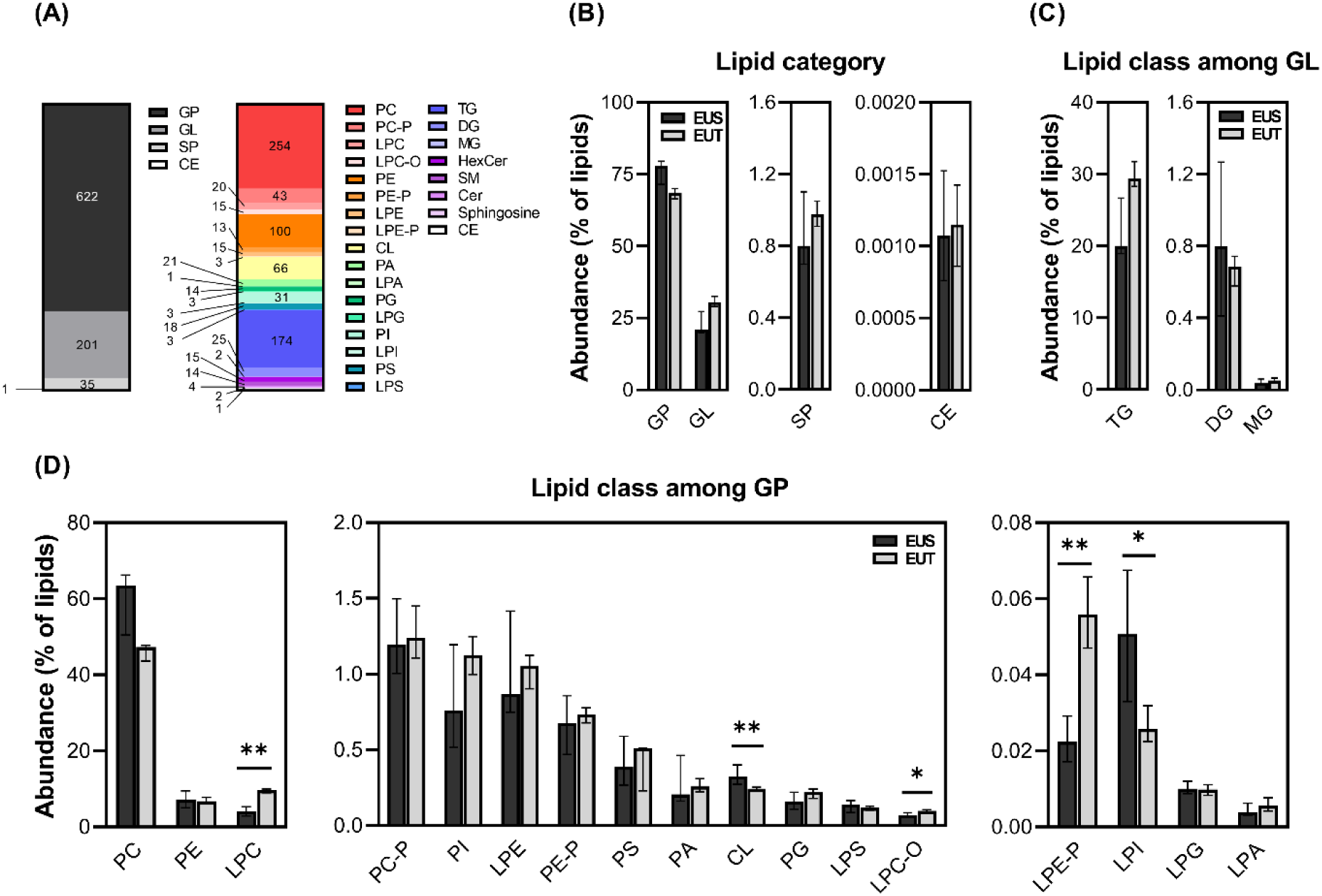
Whole-body lipidome profiles of *E. superba* and *E. triacantha*. (A) Numbers of lipid molecular species detected by LC–MS/MS across the three major lipid categories and cholesteryl esters (CE) (left) and across the 24 lipid classes within the three major categories (right). Lipid abbreviations are defined in Figure S4. (B–D) Relative abundance distributions of lipids in *E. superba* (EUS) and *E. triacantha* (EUT): (B) the three major lipid categories and CE, (C) lipid classes within glycerolipids (GL), and (D) lipid classes within glycerophospholipids (GP). Data are presented as medians with interquartile ranges (IQRs), with six biological replicates per species. Black and gray bars represent EUS and EUT, respectively. * *p* < 0.05 and ** *p* < 0.01 based on two-sided Mann– Whitney U tests.

Within the GL pool, TG accounted for more than 95% of the total GL pool in both krill species, followed by DG and MG, with no significant class-level differences observed between *E. superba* and *E. triacantha* (**Figure 2C**). In contrast, the GP pool exhibited pronounced class-level differences (**Figure 2D**). PC was the dominant GP class, accounting for approximately 80% and 67% of the total GP pool in *E. superba* and *E. triacantha*, respectively. This was followed by PE (9% and 10%, respectively) and LPC (5% and 13%, respectively). Relative to *E. triacantha*, *E. superba* exhibited significantly lower abundances of LPC, alkyl-linked LPC (LPC-O), and alkenyl-linked LPE (LPE-P), together with significantly higher abundances of CL and LPI.

To assess whether these class-level differences were also reflected at the molecular-species level, individual lipid species within the significantly different GP classes were further examined. Numerous molecular species within these classes differed significantly between the two krill species (FDR < 0.05), and their directions of change were generally consistent with the corresponding class-level patterns, with only a small number of exceptions, including CL(18:1/20:4/20:4/22:0) and LPC(18:4/0:0) (**Figure S5**). Together, whereas the GL pool showed no significant class-level differences, the GP pool exhibited class- and molecular-species-level remodeling in *E. superba*.

### Opposite Polyunsaturation Patterns in Triacylglycerols and Glycerophospholipids between *E. superba* and *E. triacantha*

To characterize the covariation patterns associated with interspecific metabolomic and lipidomic differences, Spearman correlation analysis was performed between the 113 differential metabolites and 218 differential lipid species across all 12 krill samples. Using a stringent threshold of |r| ≥ 0.9 and FDR < 0.05, lipid species belonging to PC, PE, and TG showed the largest numbers of significant cross-omics correlations with differential metabolites (**Figure 3A**). These molecular species-level covariations were observed despite the absence of significant interspecific differences in the total abundance of the PC, PE, and TG classes (**Figures 2C and 2D**), highlighting the contribution of molecular-species level variation that was not apparent from class-level abundance alone. Correlations were then examined among the differential lipid species and revealed distinct intra- and inter-class covariation patterns. Intra-class correlations were predominantly positive within TG, PC, and PE, with 328 of 328, 169 of 243, and 35 of 45 retained correlations being positive, respectively. In contrast, correlations between TG and PC were predominantly negative, with 99 of 132 retained correlations being negative, while PC–PE correlations also showed a greater number of negative than positive relationships (95 of 163 retained correlations; **Figure 3B**). These patterns motivated further examination of whether molecular differences within these major lipid classes were associated with systematic differences in lipid unsaturation.

**Figure 3.**
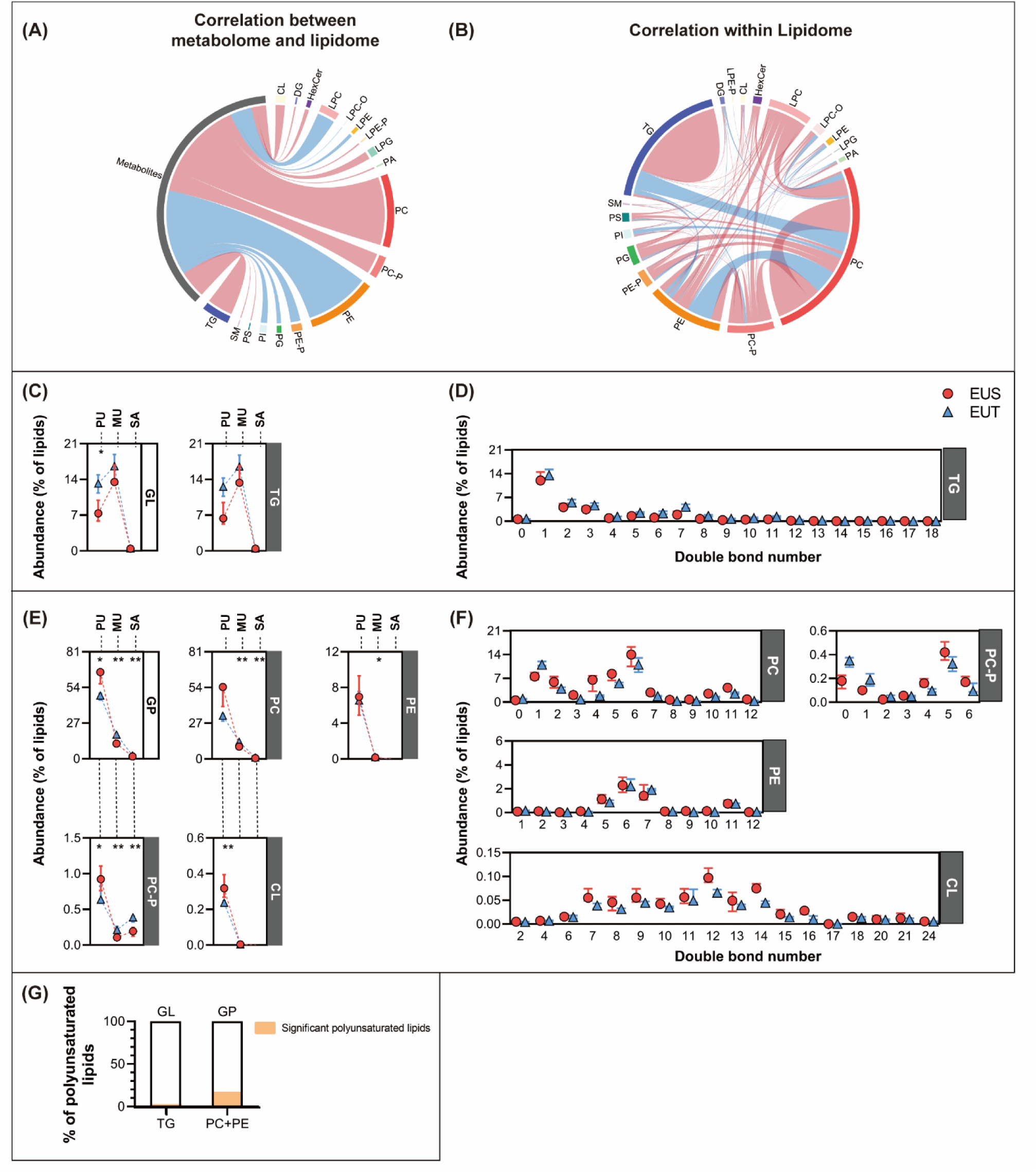
Cross-omics correlations and lipid molecular-species unsaturation patterns in *E. superba* and *E. triacantha*. (A, B) Chord diagrams showing (A) lipid–metabolite correlations and (B) lipid–lipid correlations calculated across all 12 biological samples using Spearman’s rank correlation analysis. Correlations meeting |*r*| ≥ 0.9 and FDR < 0.05 were retained and aggregated according to lipid class for visualization. In (A), metabolites are represented as a single group. Ribbon width represents the number of retained pairwise correlations between the corresponding groups, whereas ribbon color represents the direction of the mean Spearman correlation coefficient across the aggregated pairs. Red and blue ribbons indicate positive and negative mean correlations, respectively. (C, E) Relative abundances of lipid molecular species classified according to the unsaturation of their constituent fatty acyl chains: (C) GL and TG, and (E) GP and the PC, PE, PC-P, and CL. SA species contain exclusively saturated fatty acyl chains; MU species contain one or more monounsaturated fatty acyl chains but no polyunsaturated chains; and PU species contain at least one polyunsaturated fatty acyl chain. Only representative lipid classes are shown; results for the remaining lipid classes are provided in Figure S7. Data are presented as medians with interquartile ranges (IQRs), with six biological replicates per species. Red circles and blue triangles represent *E. superba* (EUS) and *E. triacantha* (EUT), respectively. * *p* < 0.05 and ** *p* < 0.01 based on two-sided Mann–Whitney U tests. (D, F) Distribution of lipid molecular-species abundances according to summed double-bond number within (D) TG and (F) PC, PE, PC-P, and CL. Data are presented as medians with interquartile ranges (IQRs). Red circles and blue triangles represent EUS and EUT, respectively. (G) Contribution of significantly different PU-classified lipid molecular species to the total PU-classified lipid abundance within the GL and GP pools of *E. superba*. For GL, the numerator comprised differential PU-classified TG species and the denominator comprised all PU-classified GL species. For GP, the numerator comprised differential PU-classified PC and PE species and the denominator comprised all PU-classified GP species. Proportions were calculated for each biological replicate, and bars represent the median across the six *E. superba* samples.

Within the GL pool, monounsaturated lipids were the most abundant in both krill species, followed by polyunsaturated and saturated lipids (**Figure 3C**). Compared with *E. triacantha*, *E. superba* exhibited significantly lower abundance of polyunsaturated GLs. Unsaturation patterns were further examined at both the class and molecular-species levels for TGs. When TGs were grouped according to unsaturation category, no significant interspecific differences were detected (**Figure 3C**). Across individual double-bond groups, TG abundance was concentrated among species with relatively low numbers of double bonds and generally declined toward higher degrees of unsaturation in both krill species (**Figure 3D**). More pronounced differences emerged at the lipid-species level: among 174 detected TG species, 34 differed significantly between the two krill species (FDR < 0.05), and all 34 were lower in *E. superba* (**Figure S6**). Of these, 33 were polyunsaturated, whereas the remaining species was the monounsaturated TG(16:0/17:0/18:1). Collectively, the differential polyunsaturated TG species accounted for only a small proportion of the total abundance of the polyunsaturated GL pool in *E. superba* (**Figure 3G**). Thus, despite the absence of a difference in total TG abundance, *E. superba* exhibited selective reduction in a subset of predominantly polyunsaturated TG molecular species.

The GP pool exhibited the opposite interspecific pattern. GPs were predominantly polyunsaturated in both krill species, followed by monounsaturated and saturated species (**Figure 3E**). Compared with *E. triacantha*, *E. superba* exhibited significantly higher abundances of polyunsaturated GPs, alongside lower abundances of saturated and monounsaturated GPs. To identify the lipid classes contributing to these differences, unsaturation patterns were analyzed across individual GP classes (**Figures 3E and S7**). Polyunsaturated PC-P and CL were significantly more abundant in *E. superba*; however, these classes together accounted for less than 2% of total GP abundance (**Figure 2D**), indicating that the overall increase in polyunsaturated GPs was primarily associated with more abundant GP classes.

Analysis of individual lipid species revealed that PC and PE contained the largest numbers of significantly different species, comprising 71 of 254 detected PC species and 34 of 100 detected PE species (FDR < 0.05), followed by PC-P with 19 of 43 species (**Figure S6**). Among the 71 differential PC species, 50 were polyunsaturated, 11 were monounsaturated, and 10 were saturated; among the 34 differential PE species, 27 were polyunsaturated and 7 were monounsaturated. *E. superba* showed selective enrichment of polyunsaturated molecular species, with higher abundance in 28 of the 50 differential polyunsaturated PC species (56.0%) and 22 of the 27 differential polyunsaturated PE species (81.5%). When grouped by double-bond number, PC and PE abundances increased toward higher degrees of unsaturation and peaked at six double bonds (**Figure 3F**). Collectively, the differential polyunsaturated PC and PE species accounted for more than one-quarter of the total abundance of the polyunsaturated GP pool in *E. superba* (**Figure 3G**), consistent with the overall enrichment of polyunsaturated GPs in *E. superba*.

Together, these results reveal contrasting molecular patterns of lipid unsaturation between the two species: there was a selective reduction of predominantly polyunsaturated TG species that occurred alongside an enrichment of polyunsaturated GPs, particularly within the PC and PE classes in *E. superba*, when compared to *E. triacantha*.

#### Distinct Distribution of Fatty Acyl Chains across Glycerophospholipid and Glycerolipid Pools

To further characterize the contrasting lipid patterns observed across GP and GL pools, fatty acyl chains were examined according to carbon-chain length, degree of unsaturation, and carbon-number parity (even-or odd-numbered). Fatty acyl abundance was estimated after normalization of total fatty acyl abundance across all lipid categories within each sample, with contributions subsequently examined according to the lipid category from which they were derived (**Figure 4, see also Figure S8**).

**Figure 4.**
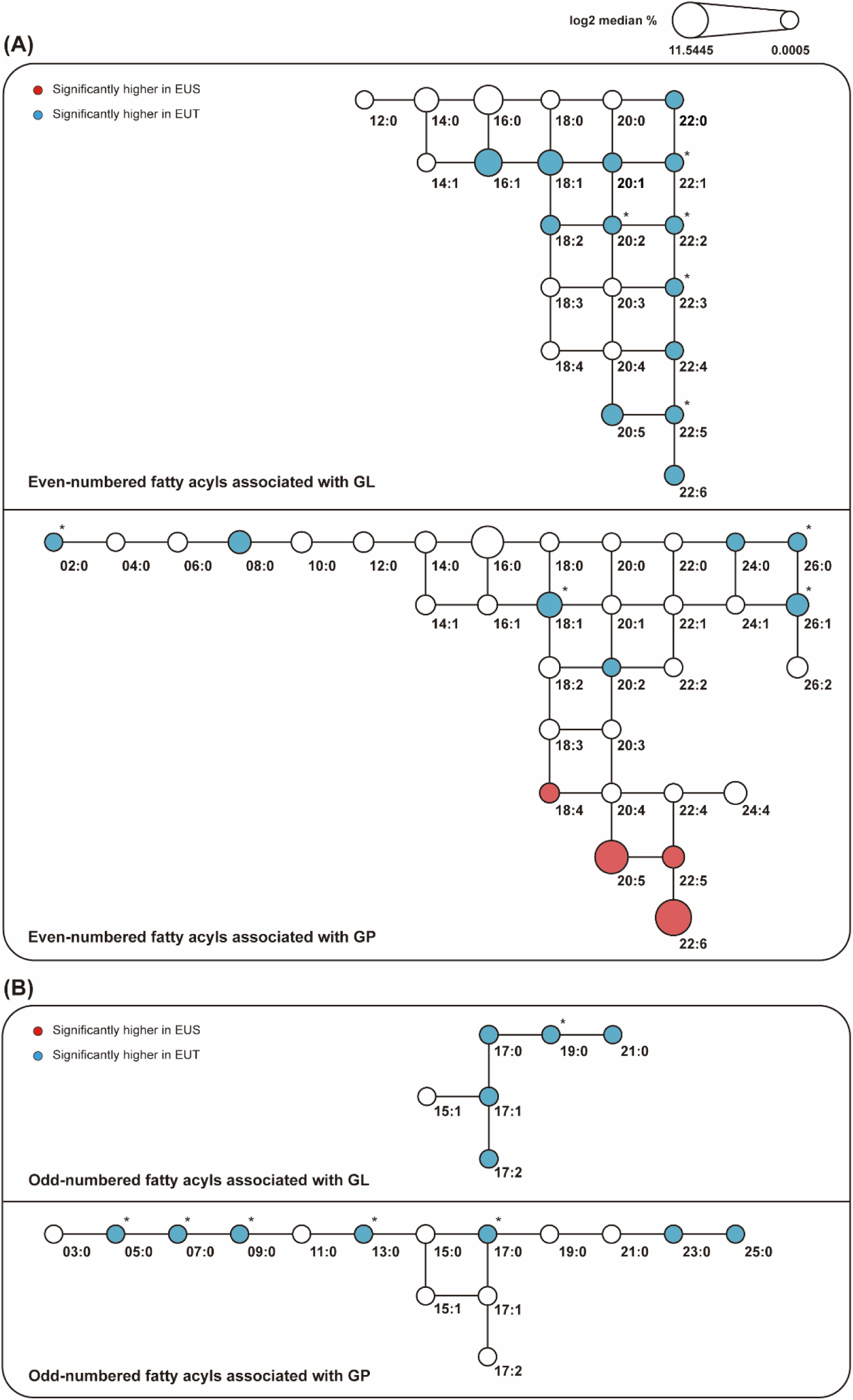
Distribution of fatty acyl chains across glycerolipid and glycerophospholipid pools. The network summarizes fatty acyl composition patterns in glycerolipid (GL) and glycerophospholipid (GP) pools and is simplified from Figure S8 to facilitate visualization. Nodes represent individual fatty acyl chains, denoted as carbon number:double-bond number. Node size represents the log2-transformed median relative abundance of each fatty acyl chain in *E. superba*. Red and blue nodes indicate fatty acyl chains with significantly higher relative abundances in *E. superba* (EUS) and *E. triacantha* (EUT), respectively, based on two-sided Mann–Whitney U tests (*p* < 0.05). Asterisks indicate fatty acyl chains that remained significant after Benjamini–Hochberg correction (FDR < 0.05). Connecting lines provide a structural framework for increasing chain length and unsaturation and do not imply specific biosynthetic pathways. (A) Even-numbered fatty acyl chains in the GL and GP pools. (B) Odd-numbered fatty acyl chains in the GL and GP pools.

Within the GL pool, *E. superba* exhibited lower abundances of multiple even-numbered fatty acyl chains relative to *E. triacantha*, with particularly pronounced reductions among C22 fatty acyl chains (**Figure 4A**). In contrast, the overall distribution of even-numbered fatty acyl chains within the GP pool was similar between the two krill species, although several individual chains differed in relative abundance. *E. superba* exhibited higher abundances of C18:4, C20:5, C22:5, and C22:6 within the GP pool, whereas C20:5, C22:5, and C22:6 were lower within the GL pool. A subset of the fatty acyl differences shown in **Figure 4** remained significant after Benjamini–Hochberg correction (FDR < 0.05). Further analysis revealed that C18:4, C20:5, C22:5, and C22:6 were predominantly associated with PC and PE species within the GP pool (**Figure S9**). Odd-numbered fatty acyl chains showed a different pattern, with *E. superba* exhibiting lower abundances across the fatty acyl chains that differed between species in both the GP and GL pools (**Figure 4B**). Although odd-numbered acyl chains represented a minor component of the overall lipidome, they were detected across most major GP and GL classes, with the exception of LPI, LPS, LPG, CL, and MG.

Despite these interspecific differences, the two krill species shared a similar overall fatty acyl framework. The detected GP pool encompassed a wide range of chain lengths, including even-numbered fatty acyls from C2 to C26 and odd-numbered fatty acyls from C3 to C25, whereas the GL pool was more restricted, primarily comprising even-numbered fatty acyls from C12 to C22 and odd-numbered fatty acyls from C17 to C21 (**Figure 4**). In terms of relative abundance, the GP fatty acyl pool was dominated by C22:6, C20:5, C18:1, and C16:0, whereas the GL fatty acyl pool was dominated by C18:1, C16:1, C16:0, and C14:0 (**Figures 4 and S8**).

Together, these results demonstrate that the two krill species share a similar overall fatty acyl framework but differ markedly in the distribution of specific fatty acyl chains across GP and GL pools. This contrast was particularly pronounced in *E. superba* for C20:5, C22:5, and C22:6 fatty acyl chains, which were more abundant within the GP pool but less abundant within the GL pool relative to *E. triacantha*.

## Discussion

### Summary

In this study, integrated metabolomic and lipidomic analyses reveal a metabolic phenotype associated with long-term cold adaptation in *E. superba*, involving highly selective lipid remodeling rather than a global increase in lipid unsaturation. C20:5, C22:5, and C22:6 fatty acyl chains showed higher relative abundances in the glycerophospholipid (GP) pool but lower relative abundances in the glycerolipid (GL) pool, alongside reduction of a subset of polyunsaturated triacylglycerol (TG) molecular species. These lipidomic differences coincided with alterations in glycerophospholipid and glycerolipid metabolism and with changes in pathways associated with carbon metabolism, nucleotide turnover, redox balance, and lipid precursor supply. Together, these patterns support an integrated metabolic phenotype in which selective fatty acyl partitioning and associated metabolic reorganization characterize long-term cold adaptation in *E. superba*.

#### A Hypothesized Metabolic Framework for Cold Adaptation in *E. superba*

Our comparative analysis captures metabolic characteristics associated with long-term cold adaptation rather than acute temperature-induced responses. Because this is an interspecific comparison, the observed biochemical differences cannot be attributed exclusively to temperature. Building on these observations, we propose a conceptual metabolic framework describing how lipid remodeling and metabolic alterations may be functionally linked in *E. superba* during long-term cold adaptation (**Figure 5**). This framework is intended to generate mechanistic hypotheses rather than establish direct causal relationships.

**Figure 5.**
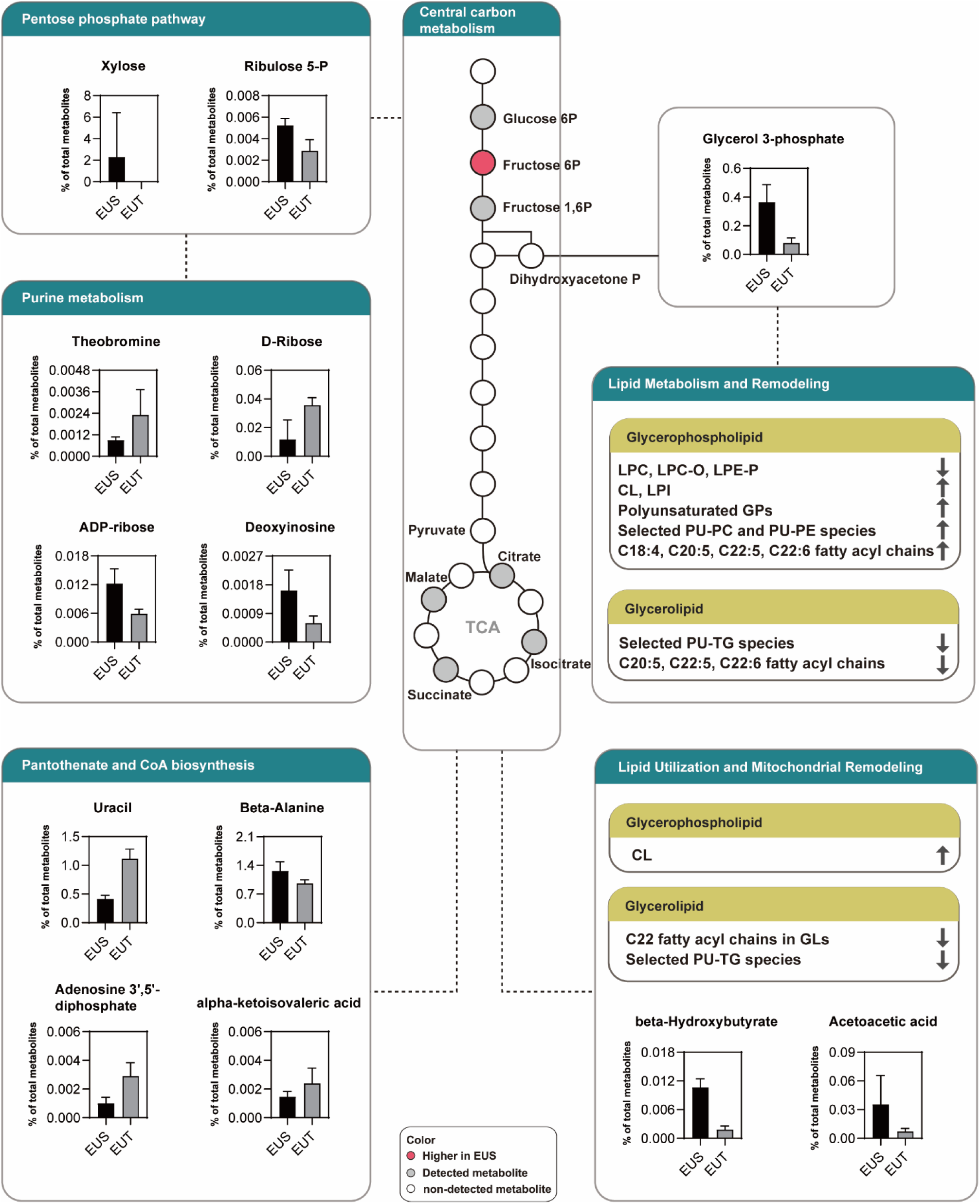
Integrated metabolic framework associated with long-term cold adaptation in *E. superba*. The schematic integrates selected metabolomic and lipidomic differences between *E. superba* (EUS) and *E. triacantha* (EUT), highlighting metabolic support for lipid remodeling, selective fatty acyl partitioning across glycerophospholipid (GP) and glycerolipid (GL) pools, and lipid utilization. The central carbon metabolism pathways are shown in full to preserve pathway context, whereas the lipid-related modules focus on features showing significant interspecific differences. Bar plots show medians with interquartile ranges (n = 6 per species), with black and gray bars representing EUS and EUT, respectively. In the lipid metabolism and remodeling and lipid utilization and mitochondrial remodeling modules, upward and downward arrows indicate higher and lower abundances, respectively, in EUS relative to EUT. Statistical significance for lipid features was determined according to the criteria used in the corresponding analyses. Dashed lines denote simplified biochemical connections. The framework does not imply direct metabolic flux or causal regulation by temperature.

### 4.1 Selective Polyunsaturated Glycerophospholipid Remodeling during Long-Term Cold Adaptation

Low temperatures challenge membrane functionality by reducing membrane fluidity and altering membrane-associated processes such as transport and signaling (Overgaard et al., 2021). Organisms commonly compensate for these effects through homeoviscous adaptation, adjusting membrane lipid composition and increasing the incorporation of unsaturated fatty acids into phospholipids to maintain appropriate membrane physical properties and function (Sinensky, 1974; Chintalapati et al., 2004).

Consistent with this framework, *E. superba* exhibited significant enrichment of polyunsaturated GPs relative to *E. triacantha* (**Figure 3E**). At the molecular-species level, this pattern was characterized by selective enrichment of polyunsaturated PC and PE species (**Figure S6**), while fatty acyl chain analysis further showed that C18:4, C20:5, C22:5, and C22:6 were predominantly associated with PC and PE within the GP pool (**Figures 4A and S9**). These patterns indicate that GP remodeling associated with long-term cold adaptation was particularly evident within PC and PE rather than being uniformly distributed across the GP pool. Recent work in congeneric species of oysters has similarly shown species-specific adjustment of glycerophospholipid unsaturation and acyl-chain length during acute temperature stress (Du et al., 2025). Head-group-specific acyl-chain remodeling has also been identified as an important feature of membrane adaptation in lipidomic studies of *Methylobacterium extorquens* and *Saccharomyces cerevisiae* during environmental or temperature change, while studies of the minimal cell JCVI-syn3B further linked the capacity for such remodeling to effective homeoviscous adaptation (Chwastek et al., 2020; Safronova et al., 2024). Biophysical experiments further showed that the effects of acyl-chain remodeling on membrane lipid packing depend on phospholipid headgroup identity, with PE showing the strongest response, PC an intermediate response, and PG a comparatively smaller response at similar levels of acyl-chain unsaturation (Chwastek et al., 2020). The selective remodeling of PC and PE observed in *E. superba* therefore suggests that long-term cold adaptation may similarly involve head-group-specific remodeling that fine-tunes membrane physical properties rather than a uniform increase in GP unsaturation. GP remodeling in *E. superba* extended beyond differences in fatty acyl unsaturation.

Among ether-linked phospholipids, LPC-O and LPE-P were significantly less abundant in *E. superba* than in *E. triacantha* (**Figure 2D**). LPC-O contains an alkyl ether linkage, whereas LPE-P contains an alkenyl ether linkage. Ether-linked phospholipids have been implicated in membrane organization and protection from lipid oxidation, with the vinyl ether bond of alkenyl ether lipids being particularly susceptible to oxidative attack (Broniec et al., 2011; Almsherqi, 2021). Together with the selective enrichment of polyunsaturated PC and PE molecular species, the lower abundance of these ether-linked lysophospholipids suggests that GP remodeling in *E. superba* involves multiple aspects of membrane lipid composition rather than a simple increase in unsaturation. Such remodeling may also be associated with altered requirements for cellular redox homeostasis, given the greater susceptibility of highly polyunsaturated lipids to oxidative damage and the changes observed in selected ether-linked phospholipid composition. In addition, LPC abundance was significantly lower in *E. superba* than in *E. triacantha* (**Figure 2D**). Because lysophospholipids serve as intermediates in phospholipid deacylation–reacylation pathways such as the Lands cycle (Lubawy et al., 2022), the reduction in LPC alongside these molecular-species changes is consistent with altered phospholipid turnover or reacylation. Lower LPC abundance may therefore reflect changes in lysophospholipid metabolism associated with the maintenance of specific PC molecular species in *E. superba*.

Intriguingly, the enrichment of polyunsaturated GPs was accompanied by opposite patterns of C20:5, C22:5, and C22:6 fatty acyl abundance between the GP and GL pools, with higher relative abundances in GPs but lower relative abundances in GLs in *E. superba* (**Figure 4A**). Given that TG accounted for more than 95% of the GL pool and that predominantly polyunsaturated TG molecular species were selectively reduced (**Figures 2C and S6**), these observations together strengthen the contrast between membrane-associated and storage lipid pools. We interpret this opposite pattern as evidence of selective fatty acyl partitioning across GP and GL pools. This interpretation does not imply direct transfer of fatty acyl chains from TGs to GPs; rather, the contrasting profiles could arise from differences in fatty acid synthesis, activation, incorporation, remodeling, storage, or utilization. Such selectivity is biochemically plausible because enzymes involved in lipid synthesis and remodeling often exhibit substrate preferences. *E. superba* has previously been reported to retain substantial levels of polyunsaturated fatty acids despite variation in dietary polyunsaturated fatty acid composition, and feeding experiments have suggested some capacity for endogenous modification of dietary fatty acids (Virtue et al., 1993b). In other metazoans, long-chain acyl-CoA synthetases (ACSLs) display differential substrate preferences, with ACSL4 favoring C20:4(n-6) and C20:5(n-3), whereas ACSL6 exhibits comparatively high activity toward C22:6(n-3) (Kang et al., 1997; Marszalek et al., 2005). Lysophospholipid acyltransferases also contribute to the selective generation of phospholipid molecular species (Yamashita et al., 2014), providing a plausible biochemical basis for selective incorporation of polyunsaturated fatty acids into GPs.

Although whole-body lipidomic analysis cannot resolve tissue- or organelle-specific membrane compositions, the consistency of these molecular patterns supports extensive remodeling of the GP pool associated with long-term adaptation to the Antarctic cold environment. Together, the convergence of lipid-class, molecular-species, unsaturation, and fatty-acyl analyses suggests that selective polyunsaturated GP remodeling is a prominent feature of membrane adaptation in *E. superba*.

### 4.2 Metabolic Reorganization Accompanying Lipid Remodeling during Long-Term Cold Adaptation

The extensive lipid remodeling observed in *E. superba* was accompanied by concurrent changes in pathways associated with carbon metabolism, nucleotide turnover, redox balance, and lipid precursor supply (**Figure 5**). Rather than representing isolated metabolic differences, these alterations suggest a reorganization of cellular metabolism that may provide the biosynthetic substrates and redox support needed to sustain lipid remodeling and cellular homeostasis under long-term cold conditions.

One prominent component of this metabolic reorganization was the enrichment of metabolites associated with the pentose phosphate pathway (PPP), together with significant enrichment of pentose and glucuronate interconversions and purine metabolism (**Figure 1B**). Cold-responsive changes involving nucleotide metabolism and redox-associated processes have also been reported in transcriptomic and metabolomic studies of fish, crustaceans, and yeast (Chen et al., 2008; Jiao et al., 2020; Ren et al., 2020; Ronges et al., 2012; Wu et al., 2019; Touchette et al., 2022), suggesting that sustained adjustment of nucleotide and redox metabolism may represent an important component of cold adaptation. These adjustments may help compensate for reduced biochemical reaction rates that constrain macromolecular synthesis, repair, and turnover at low temperature (Peck, 2016). Antarctic fishes, for example, maintain elevated RNA-to-protein ratios despite relatively low translational rates (Fraser et al., 2022), while constitutive expression of molecular chaperones such as HSP70 has been reported in *E. superba* (Clark and Peck, 2009; Toullec et al., 2020). These observations suggest a role for enriched nucleotide-associated pathways in maintaining macromolecular synthesis, repair, and turnover under cold conditions.

Moreover, the PPP provides a direct biochemical link between central carbon metabolism and cellular redox homeostasis through the generation of NADPH (TeSlaa et al., 2023). NADPH supplies reducing equivalents required for fatty acid biosynthesis and desaturation while also supporting glutathione-and thioredoxin-dependent antioxidant systems (Stincone et al., 2015). This function may be particularly relevant in *E. superba*, where the enrichment of highly polyunsaturated GP species may increase the susceptibility of membrane lipids to oxidative damage. Enrichment of PPP-associated metabolism is therefore consistent with metabolic adjustment The observed changes in PPP-associated metabolites are therefore consistent with a metabolic context that could support both reductive biosynthesis and redox homeostasis.

Changes in glycerol-3-phosphate metabolism provide an additional connection between central carbon metabolism and lipid biosynthesis. *E. superba* exhibited significantly higher glycerol-3-phosphate abundance (**Figures 1C and 5**), consistent with altered abundance of a central precursor for both GP and GL synthesis. Derived from the glycolytic intermediate dihydroxyacetone phosphate (DHAP), glycerol-3-phosphate represents a key branch point through which carbohydrate-derived carbon can be directed toward lipid biosynthesis (Madiraju et al., 2026). Moreover, the reversible conversion between DHAP and glycerol-3-phosphate is coupled to cellular NADH/NAD⁺ balance (Liu et al., 2021), providing an additional link between lipid precursor metabolism and cellular redox state. The higher abundances of NAD and nicotinamide in *E. superba* (**Figure 1D**) are consistent with this metabolic adjustment.

Taken together, changes in nucleotide-associated metabolism, the PPP, glycerol-3-phosphate, and NAD/nicotinamide abundance place lipid remodeling in *E. superba* within a metabolic context involving biosynthetic maintenance, lipid-precursor metabolism, and redox-associated processes. These adjustments may help sustain the remodeled lipid state associated with long-term cold adaptation.

### 4.3 Lipid Utilization and Mitochondrial Membrane Remodeling during Long-Term Cold Adaptation

While the contrasting distribution of C20:5, C22:5, and C22:6 between the GP and GL pools supports selective fatty acyl partitioning, the reduction of C22 fatty acyl chains within GLs in *E. superba* suggests an additional metabolic dimension involving storage-lipid utilization (**Figure 4A**). Because TG accounted for more than 95% of the GL pool (**Figure 2C**), the selective reduction of predominantly polyunsaturated TG molecular species further supports alteration of the major neutral storage-lipid pool in *E. superba*. This pattern may reflect differential mobilization or utilization of selected polyunsaturated fatty acids, potentially contributing to the energetic demands associated with cellular maintenance, biosynthetic activity, and redox homeostasis under long-term cold conditions. Selective utilization of polyunsaturated fatty acids has previously been reported during starvation in *E. superba* (Virtue et al., 1993a), providing physiological precedent for selective utilization of these lipid reserves under increased energetic demand.

The metabolomic profile provides additional evidence for altered fatty-acid-associated metabolism. Arachidonic acid and α-linolenic acid metabolism were significantly enriched, while the ketone bodies acetoacetate and β-hydroxybutyrate were elevated in *E. superba* (**Figures 1B and 5**). Together with the reduction of polyunsaturated TG species, these changes are consistent with altered utilization or turnover of fatty-acid-derived carbon. One possible pathway contributing to altered fatty acid utilization is peroxisomal β-oxidation, which contributes to the shortening of very-long-chain and selected highly unsaturated fatty acids before further mitochondrial oxidation (Hiltunen et al., 1986). Notably, peroxisomal β-oxidation contributes approximately 30% of hepatic fatty acid oxidation in the Antarctic teleost *Notothenia gibberifrons* (Crockett and Sidell, 1993), providing physiological precedent for substantial peroxisomal fatty-acid oxidation in a polar ectotherm. These observations are also consistent with the view of peroxisomes as metabolic hubs integrating lipid metabolism with environmental stress responses (He et al., 2021).

Mitochondrial membrane remodeling may complement these changes in lipid utilization. *E. superba* exhibited significantly elevated cardiolipin (CL) abundance (**Figure 2D**), a signature phospholipid of the mitochondrial inner membrane that supports cristae organization, respiratory-chain supercomplexes, and oxidative phosphorylation (Falabella et al., 2021). Cold-associated increases or remodeling of CL have been reported across organisms ranging from unicellular systems to thermogenic mammalian tissues (Chung et al., 2018; Lynes et al., 2018; Safronova et al., 2024; Sustarsic et al., 2018), suggesting that mitochondrial membrane remodeling represents a recurrent response associated with cold exposure and adaptation. The elevated CL abundance observed in *E. superba* may therefore represent an adjustment of mitochondrial membrane organization associated with long-term cold adaptation. Interestingly, phosphatidylglycerol (PG), the immediate precursor of CL biosynthesis, did not show a corresponding increase (**Figure 2D**). This differs from transient precursor changes reported during acute cold-induced CL remodeling (Lynes et al., 2018) and may indicate that the elevated CL abundance reflects a more persistent mitochondrial membrane state rather than a transient precursor accumulation associated with acute cold exposure.

Odd-chain fatty acid metabolism may provide an additional link between lipid utilization and central carbon metabolism because β-oxidation of odd-chain fatty acids generates propionyl-CoA, which can enter the TCA cycle through succinyl-CoA formation (Inigo et al., 2021). The lower abundance of several odd-numbered fatty acyls in *E. superba* (**Figure 4B**) is therefore consistent with altered odd-chain fatty acid metabolism, although their specific metabolic fate cannot be resolved from the present data. In parallel, elevated β-alanine and enrichment of pantothenate and CoA biosynthesis (**Figures 1B and 5**) suggest adjustment of CoA-related metabolism. Because CoA is required for fatty acid activation, β-oxidation, ketogenesis, and TCA-cycle metabolism (Leonardi and Jackowski, 2007), these changes may facilitate sustained lipid turnover during long-term cold adaptation.

Together, the reduction of C22 fatty acyl chains within the GL pool and of polyunsaturated TG molecular species, along with altered fatty-acid-associated metabolism, elevated ketone bodies, and increased CL, suggests reorganization of lipid utilization and mitochondrial metabolism in *E. superba*. We therefore propose that selective fatty acyl partitioning across GP and GL pools is accompanied by differential utilization of storage lipids, linking membrane adaptation with metabolic reorganization that may support cellular homeostasis under long-term cold conditions.

### 4.4 Ecological Implications of Lipid Allocation Strategies in Euphausiids

The integrated metabolic framework proposed here suggests that long-term adaptation of *E. superba* to the Antarctic environment is associated with more than differences in total lipid abundance or overall unsaturation. Instead, the contrasting distribution of polyunsaturated fatty acyl chains across membrane-associated GP and storage-lipid pools points to an additional dimension of lipid organization: the selective association of particular fatty acyl chains with lipid pools that fulfill distinct structural and metabolic functions.

Lipid-storage strategies differ considerably among euphausiids occupying contrasting thermal and nutritional environments. Wax esters constitute major storage lipids in several high-latitude species, including *E. crystallorophias* (Bottino, 1975; Kattner and Hagen, 1998; Ju and Harvey, 2004) and *Thysanoessa macrura* (Hagen and Kattner, 1998), whereas *T. inermis* uses a more mixed storage strategy involving wax esters, TGs, and glycerophospholipids (Saether et al., 1986). Wax esters have been associated with prolonged energy storage and overwintering under strongly seasonal food availability (Bottino, 1975; Hagen and Kattner, 1998; Kattner and Hagen, 1998; Ju and Harvey, 2004). By contrast, TGs represent the dominant neutral storage lipid in species such as *E. vallentini* and *Meganyctiphanes norvegica*, whereas *T. raschii* exhibits a mixed storage pattern in which TGs and glycerophospholipids make substantial contributions (Mayzaud et al., 2003; Saether et al., 1986). These interspecific differences highlight substantial ecological diversity in euphausiid lipid-storage strategies and their associations with habitat, trophic conditions, and life-history demands.

Phospholipids may add further complexity to this classical division between wax-ester- and TG-dominated storage strategies. Across euphausiid species, GP pools vary with season, nutritional condition, developmental stage, and reproductive investment rather than behaving as completely static structural components (Hagen et al., 2001; Mayzaud et al., 2003; Saether et al., 1986). Recent high-resolution analyses of *E. superba* further revealed seasonal remodeling of intact phospholipid molecular composition within the digestive system (Heyen et al., 2023), reinforcing the view that phospholipid pools can respond dynamically to physiological state. At the whole-organism level, seasonal and starvation studies likewise demonstrate dynamic adjustment of lipid reserves in response to physiological state (Meyer et al., 2010; Auerswald et al., 2015). Histological observations in *E. superba* have additionally shown close spatial associations between neutral and phospholipid-rich lipid deposits within tissues involved in storage and metabolism (Saether et al., 1985). Recent phospholipidomic studies of marine ectotherms further indicate that membrane-lipid responses to environmental stress can vary with population and ecological origin, suggesting that thermal adaptation may involve context-dependent lipid organization rather than a universal shift in overall unsaturation (Feugere et al., 2025). Together, these observations suggest that euphausiid lipid allocation is not defined solely by the use of phospholipids as structural components and neutral lipids as energy stores, but also by the dynamic regulation of these lipid pools across physiological and environmental contexts.

Within this ecological context, the present study identifies molecular-level fatty acyl partitioning as an additional dimension of euphausiid lipid adaptation. Preferential association of polyunsaturated fatty acyl chains with GP rather than the major neutral storage pool may help maintain membrane function while linking membrane remodeling with altered storage-lipid utilization. Integrating these lipidomic patterns with metabolomic changes further places this molecular organization within an integrated metabolic context associated with long-term cold adaptation. Future comparative and functional studies will be needed to determine the extent to which this form of lipid organization occurs across euphausiids and to test the mechanisms underlying its plasticity under changing thermal environments.

## Aknowlegments

JYT benefited from funding provided by Institut Paul Emile Victor (IPEV) (KREVET program) and also from the“Région Bretagne” (SAD-1–DRAKAR program) for Antarctic campains. JYT would like to thank all the IPEV staff at the base Dumont d’Urville and Astrolabe’s crew for their help during Antarctic campaigns in Terre Adélie. JYT and CYL would also like to thank Philippe Koubbi for welcoming them into the REPCCOAI program, as well as IPEV for its logistical support, and, of course, the Marion Dufresne’s crew for their invaluable assistance. CYL was financially supported by the National Science and Technology Council (NSTC), Taiwan through grants (112-2311-B-018-001; 113-2311-B-018-001). Special thanks were due to Dr. Chi-Fong Chang (Genomic Research Center, Academia Sinica), who provided excellent technical steering to SHS.

## Supplemental information

**Figure S1.**
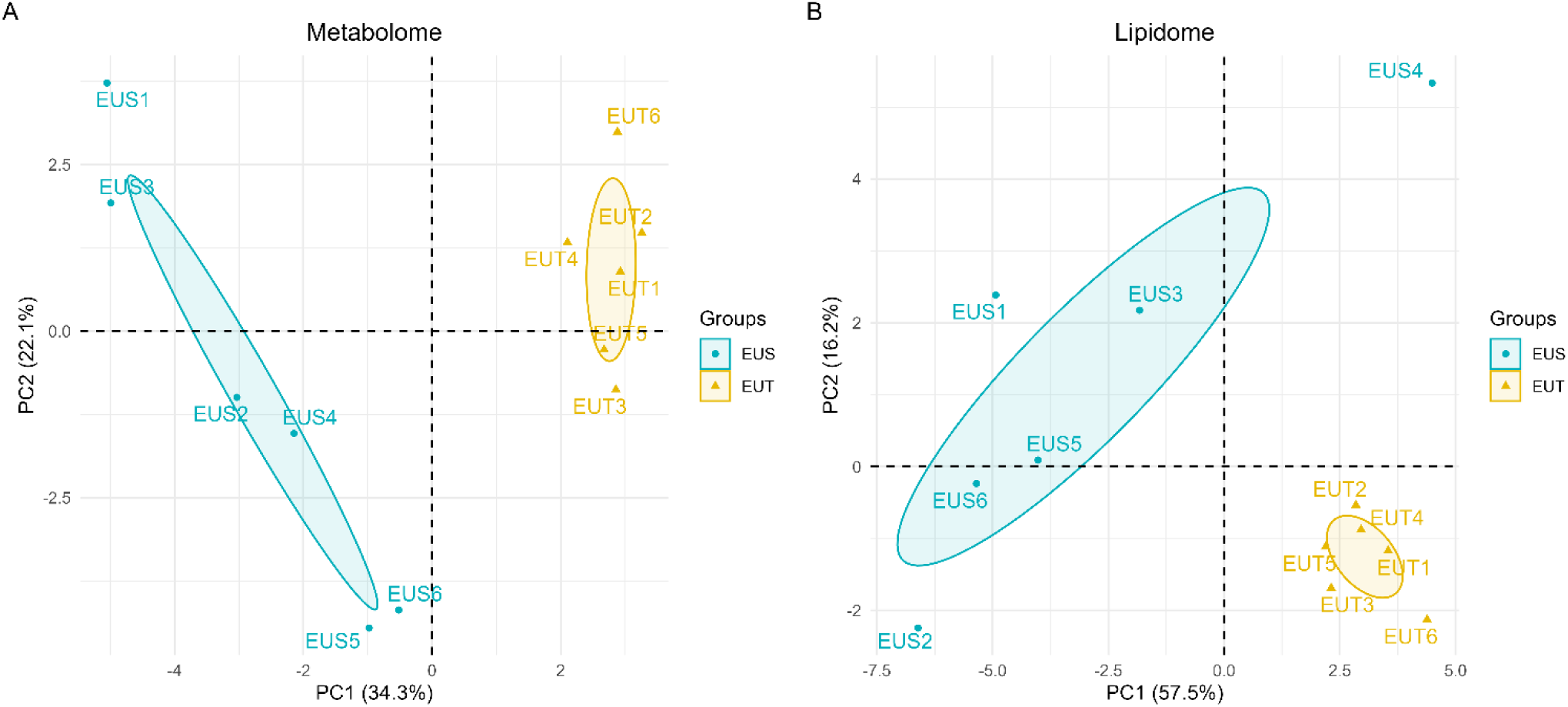
Principal component analysis (PCA) of the metabolomic and lipidomic profiles of *E. superba* and *E. triacantha*. Score plots showing (A) the whole-body metabolome and (B) lipidome profiles of *E. superba* (EUS; blue dots) and *E. triacantha* (EUT; yellow triangles). Each point represents one biological replicate (n = 6 per species). The percentages of total variance explained by PC1 and PC2 are indicated on the axes. Ellipses denote 95% confidence regions for each species. Differences in multivariate composition between species were significant by PERMANOVA for both the metabolome (R^2^ = 0.316*, p* = 0.002) and lipidome (R^2^ = 0.386, *p* = 0.013). No significant differences in multivariate dispersion were detected between species for either the metabolome (PERMDISP, *p* = 0.116) or lipidome (PERMDISP, *p* = 0.195).

**Figure S2.**

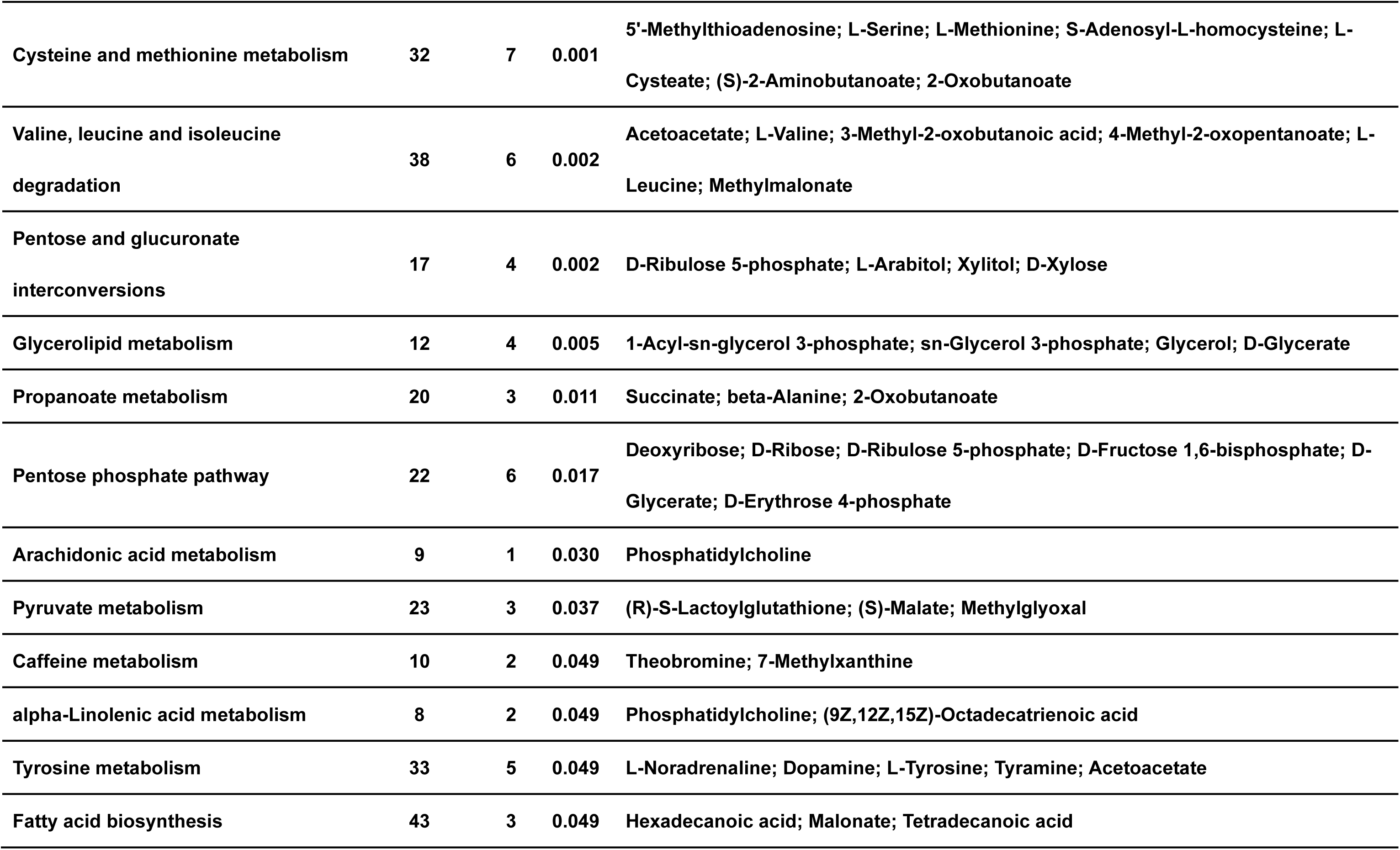
Significantly enriched metabolic pathways identified in the metabolomic analysis.

**Figure S3.**
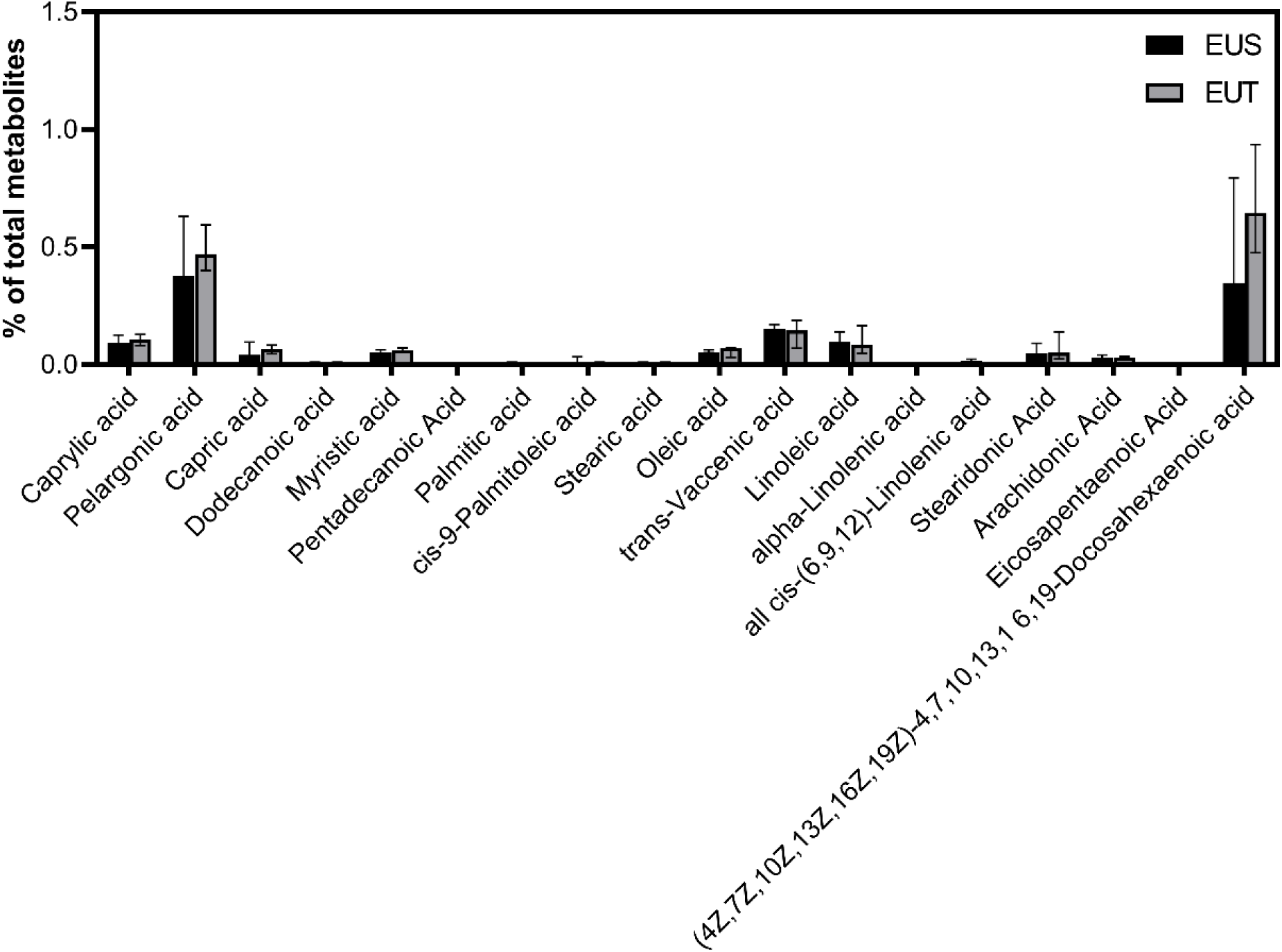
Relative abundance of free fatty acids detected in the metabolomic dataset. Individual free fatty acids (FFAs) are expressed as percentages of the total detected metabolite abundance to provide an indication of potential lipid degradation during sample processing. Individual FFA species each remained below 1% of total metabolite abundance in both *E. superba* (EUS; black bars) and *E. triacantha* (EUT; gray bars), indicating limited accumulation of free fatty acids. Data are presented as medians with interquartile ranges (IQRs), with six biological replicates per species.

**Figure S4.**
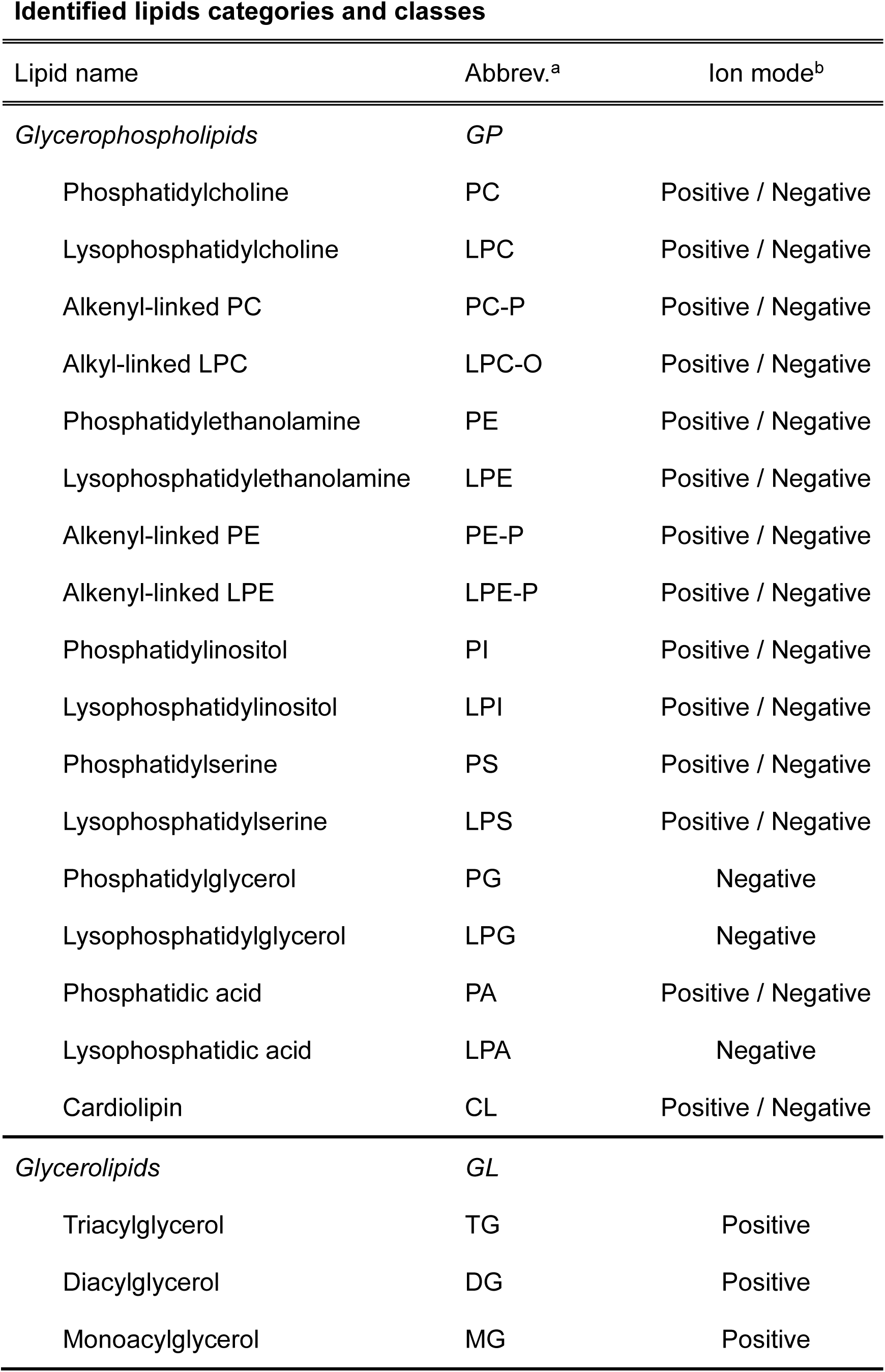
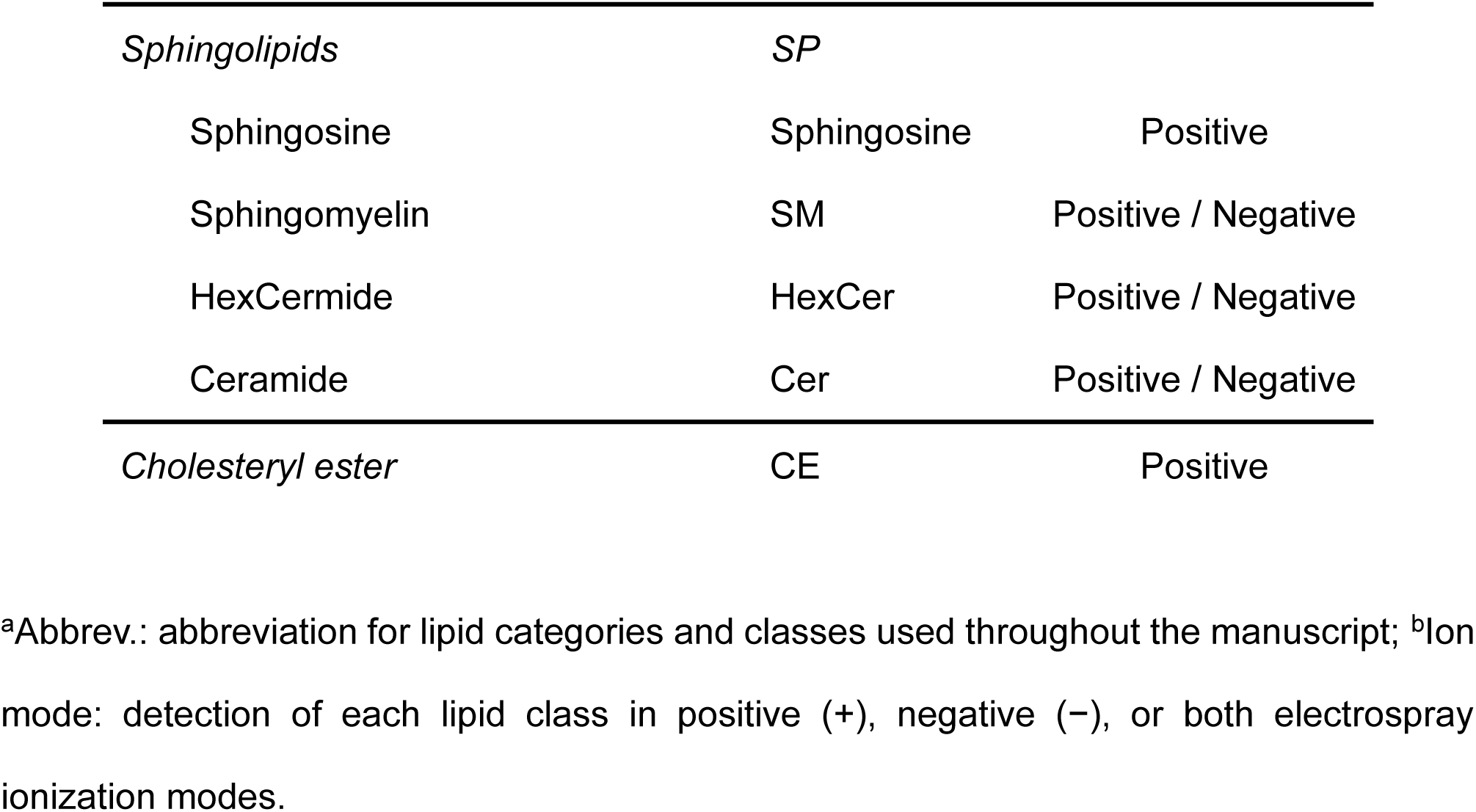
Lipid classification, abbreviations, and ionization modes used in the lipidomic analysis.

**Figure S5.**
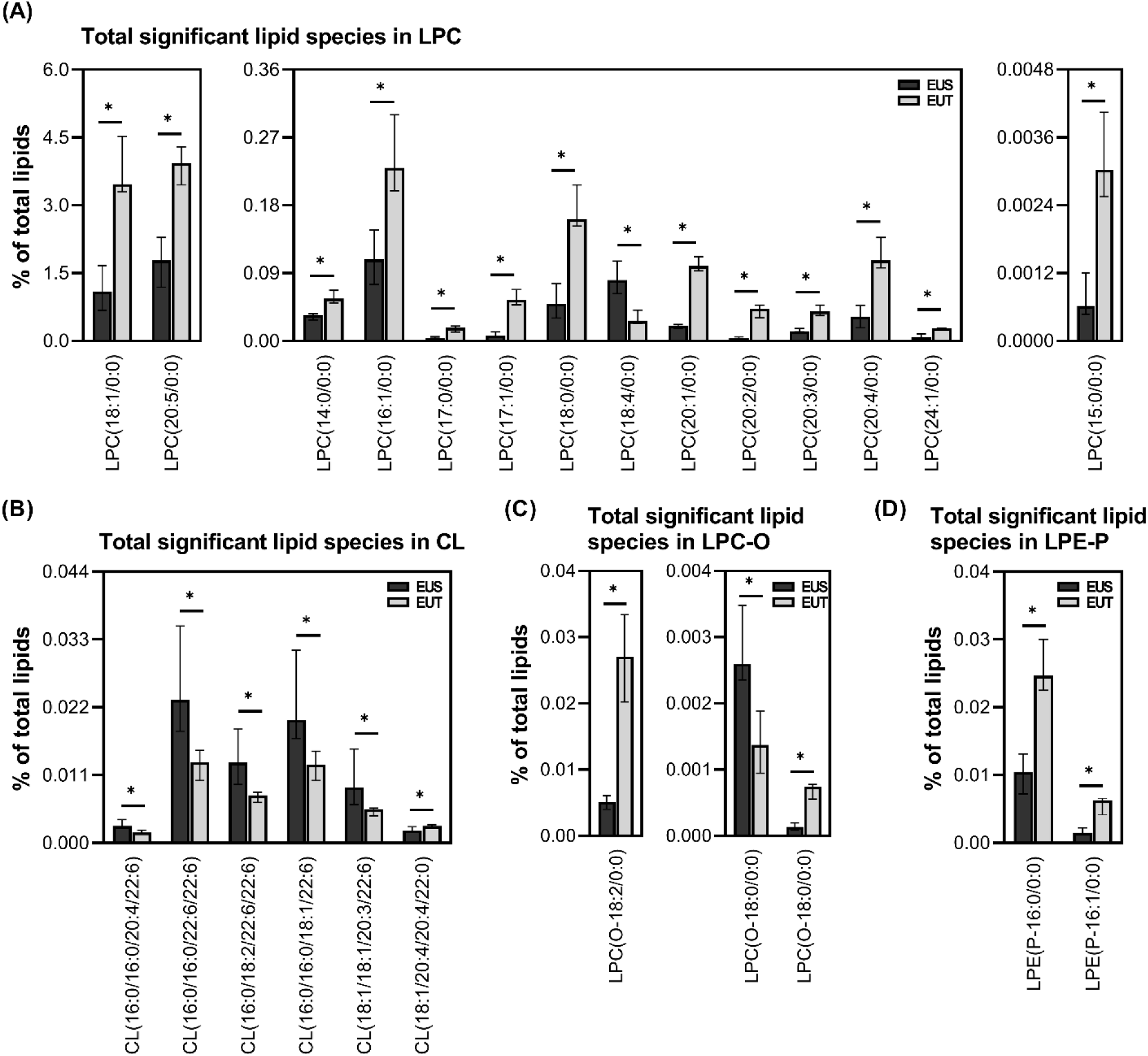
Differential lipid molecular species within selected lipid classes. Relative abundances of individual lipid molecular species within the LPC, CL, LPC-O, and LPE-P that differed significantly between *E. superba* (EUS; black bars) and *E. triacantha* (EUT; gray bars). These lipid species correspond to class-level differences shown in Figure 2D. LPI is not shown because no individual LPI species met the significance criterion. Data are expressed as percentages of total lipid abundance and presented as medians with interquartile ranges (IQRs), with six biological replicates per species. Statistical significance was assessed using two-sided Mann–Whitney U tests with Benjamini–Hochberg correction; *FDR < 0.05 and **FDR < 0.01.

**Figure S6.**

Numbers of differential lipid molecular species across lipid classes and unsaturation categories. The table summarizes lipid molecular species with significantly higher relative abundance in *E. superba* (EUS) or *E. triacantha* (EUT) within each lipid class. Differential species were identified using two-sided Mann–Whitney U tests with Benjamini–Hochberg correction (FDR < 0.05). Lipid molecular species were classified as saturated (SA), monounsaturated (MU), or polyunsaturated (PU) according to the unsaturation of their constituent fatty acyl chains, as defined in the Methods. The “Significant numbers” column indicates the total number of differential species within each lipid class, and the “Total numbers” column indicates the total number of detected lipid species in that class.

**Figure S7.**
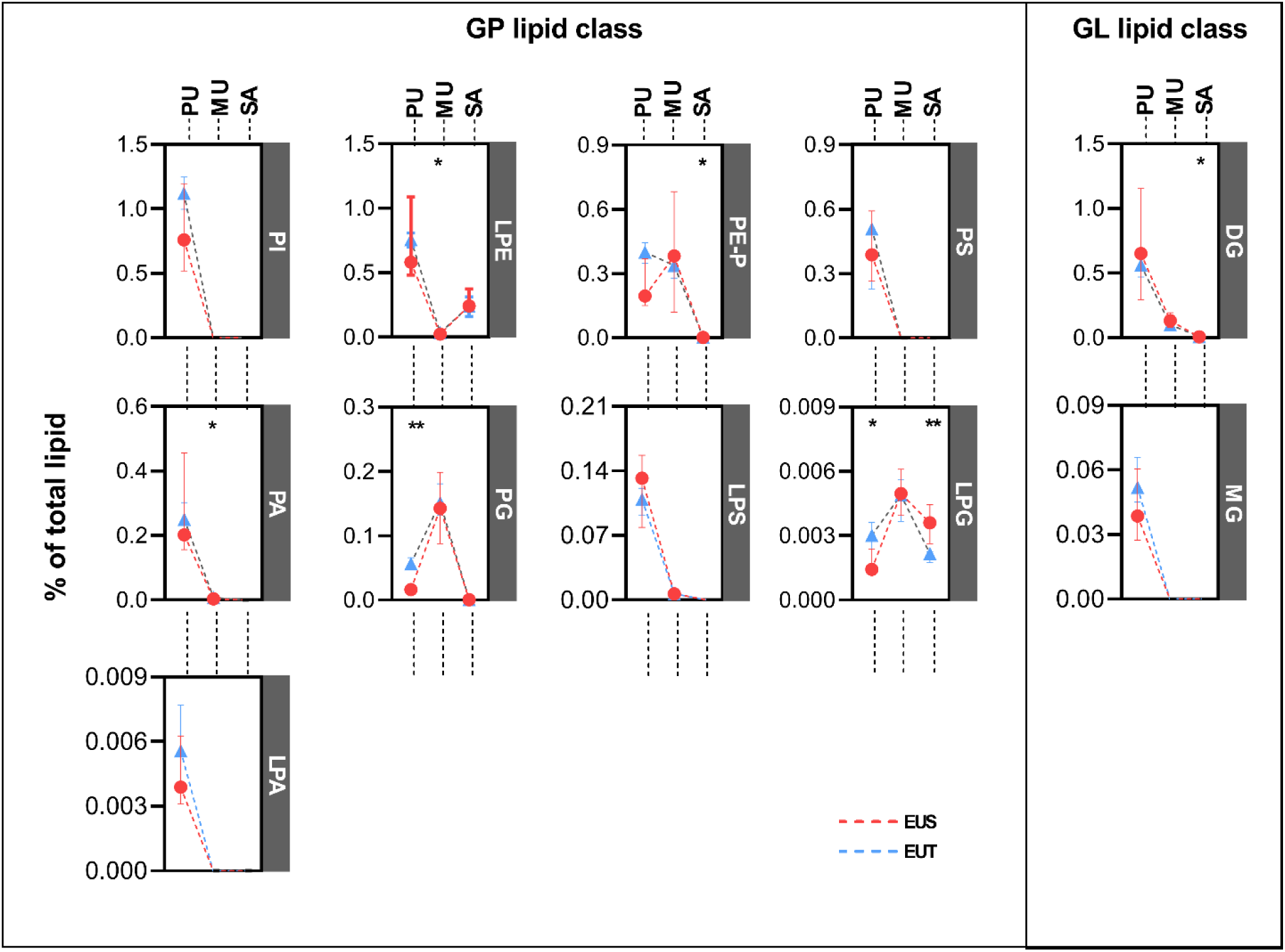
Unsaturation-category profiles across selected lipid classes. Relative abundances of lipid molecular species classified into saturated (SA), monounsaturated (MU), and polyunsaturated (PU) categories within lipid classes not shown in Figure 3. Classification was based on the unsaturation of constituent fatty acyl chains, as defined in the Materials and Methods. Data are organized by glycerophospholipid (GP) and glycerolipid (GL) categories and presented as medians with interquartile ranges (IQRs), with six biological replicates per species. *E. superba* (EUS) is shown as red circles/lines and *E. triacantha* (EUT) as blue triangles/lines. Statistical significance was determined using two-sided Mann–Whitney U tests; \**p* < 0.05 and \*\**p* < 0.01.

**Figure S8.**
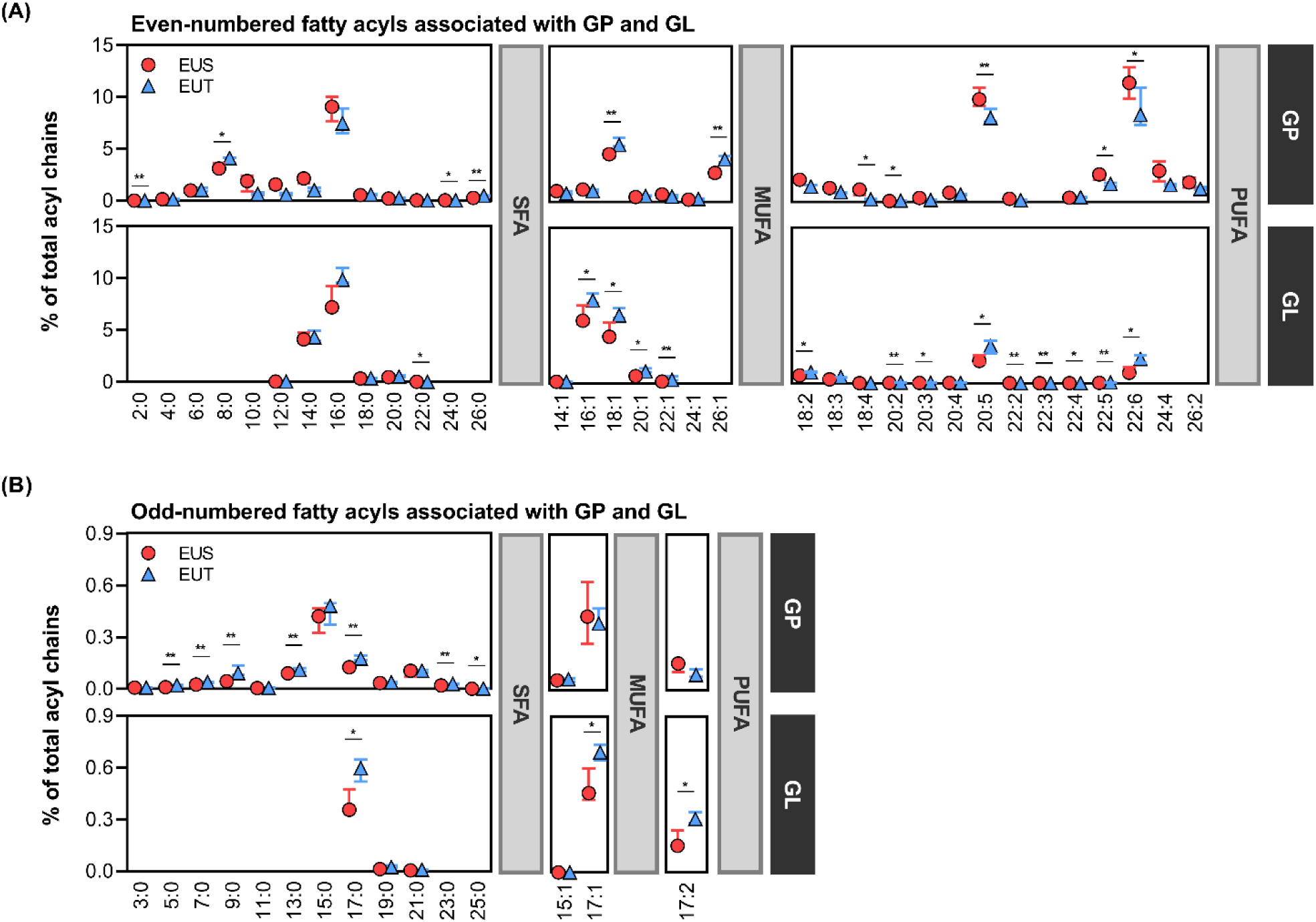
Distribution of individual fatty acyl chains within glycerophospholipid and glycerolipid pools. Relative abundances of individual fatty acyl chains derived from glycerophospholipid (GP) and glycerolipid (GL) pools in *E. superba* (EUS; red circles) and *E. triacantha* (EUT; blue triangles). Fatty acyl chains are grouped into (A) even-numbered and (B) odd-numbered fatty acyl chains and further organized according to unsaturation category. For each fatty acyl chain, the plotted abundance represents its proportional contribution after total fatty acyl abundance across all lipid categories within each sample was normalized to 100%. GP- and GL-associated contributions are displayed separately according to the lipid category from which they were derived. Data are presented as medians with interquartile ranges (IQRs), with six biological replicates per species. Statistical significance was determined using two-sided Mann–Whitney U tests; \**p* < 0.05 and \*\**p* < 0.01. Fatty acyl chains remaining significant after Benjamini–Hochberg correction (FDR < 0.05) are indicated in Figure 4. Abbreviations: SFA, saturated fatty acyl chains; MUFA, monounsaturated fatty acyl chains; PUFA, polyunsaturated fatty acyl chains.

**Figure S9.**
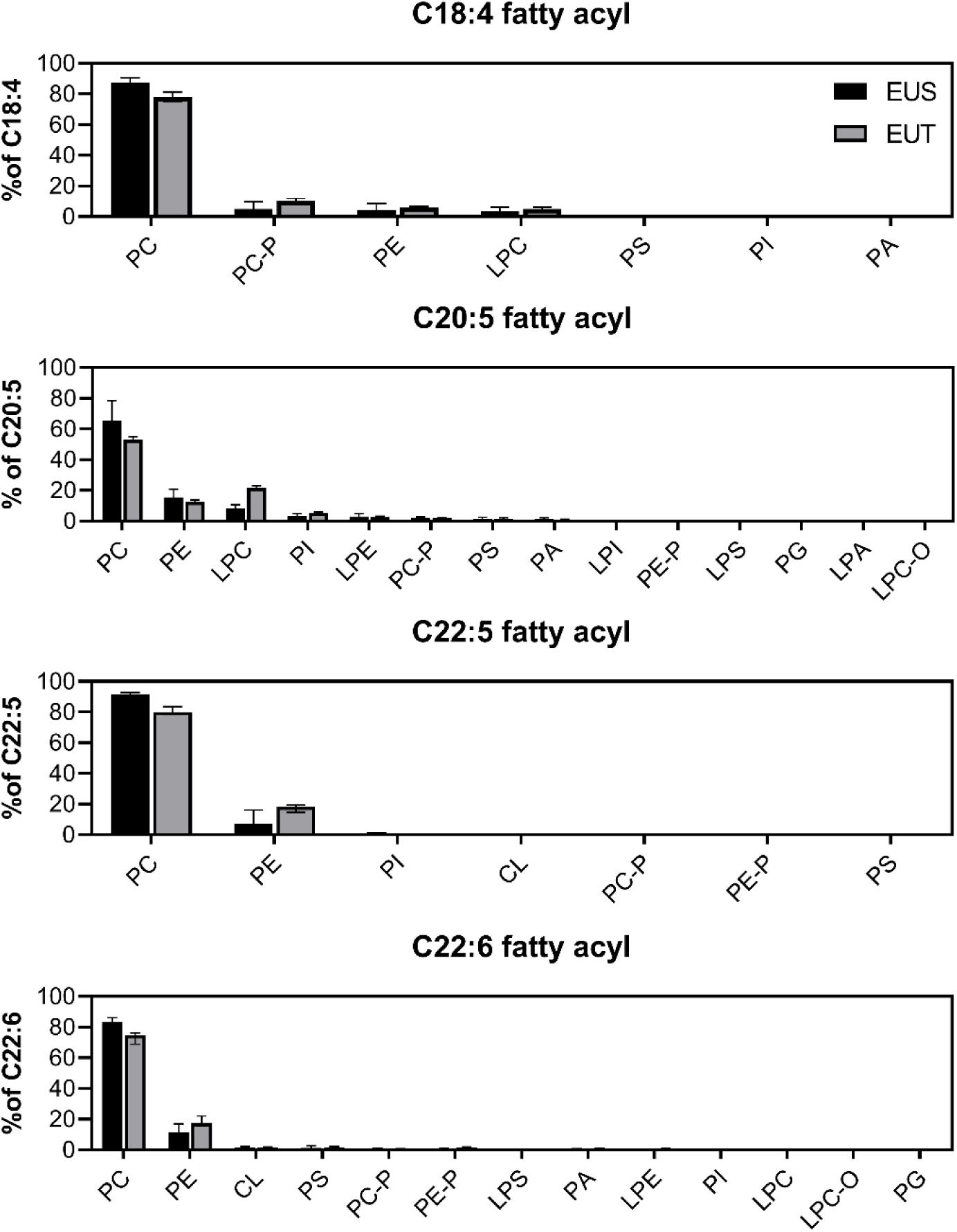
Distribution of selected polyunsaturated fatty acyl chains across glycerophospholipid classes. Each panel shows the percentage contribution of a selected polyunsaturated fatty acyl chain (C18:4, C20:5, C22:5, and C22:6) across glycerophospholipid classes. For each fatty acyl chain, its abundance was normalized across all identified glycerophospholipid classes within each sample to visualize its relative distribution among lipid classes. Data are presented as medians with interquartile ranges (IQRs), with six biological replicates per species. Black and gray bars represent *E. superba* (EUS) and *E. triacantha* (EUT), respectively.

## Notes

### Competing Interest Statement

The authors have declared no competing interest.

## References

Almsherqi Z. A. (2021). Potential Role of Plasmalogens in the Modulation of Biomembrane Morphology. Frontiers in cell and developmental biology, 9, 673917.

Atkinson, A., Siegel, V., Pakhomov, E. A., Jessopp, M. J., & Loeb, V. (2009). A re-appraisal of the total biomass and annual production of Antarctic krill. Deep Sea Research Part I: Oceanographic Research Papers, 56(5), 727–740.

Auerswald L., Pape C., Stübing D., Lopata A., Meyer B. (2009). Effect of short-term starvation of adult Antarctic krill, *Euphausia superba*, at the onset of summer. Journal of Experimental Marine Biology and Ecology, 381(*1*), 47–56.

Auerswald, L., Meyer, B., Teschke, M., Hagen, W. and Kawaguchi, S. (2015). Physiological response of adult Antarctic krill, *Euphausia superba*, to long-term starvation. Polar Biology, 38, 763–780.

Benjamini, Y., & Hochberg, Y. (1995). Controlling the False Discovery Rate: A Practical and Powerful Approach to Multiple Testing. Journal of the Royal Statistical Society. Series B (Methodological), 57(1), 289–300.

Bottino N. R. (1975). Lipid composition of two species of Antarctic krill: Euphausia superba and E. crystallorophias. Comparative biochemistry and physiology. B, Comparative biochemistry, 50(3), 479–484.

Broniec, A., Klosinski, R., Pawlak, A., Wrona-Krol, M., Thompson, D., & Sarna, T. (2011). Interactions of plasmalogens and their diacyl analogs with singlet oxygen in selected model systems. Free radical biology & medicine, 50(7), 892–898.

Cavan, E. L., Belcher, A., Atkinson, A., Hill, S. L., Kawaguchi, S., McCormack, S., Meyer, B., Nicol, S., Ratnarajah, L., Schmidt, K., Steinberg, D. K., Tarling, G. A., & Boyd, P. W. (2019). The importance of Antarctic krill in biogeochemical cycles. Nature communications, 10(1), 4742.

Chen, Z., Cheng, C. H., Zhang, J., Cao, L., Chen, L., Zhou, L., Jin, Y., Ye, H., Deng, C., Dai, Z., Xu, Q., Hu, P., Sun, S., Shen, Y., & Chen, L. (2008). Transcriptomic and genomic evolution under constant cold in Antarctic notothenioid fish. Proceedings of the National Academy of Sciences of the United States of America, 105(35), 12944–12949.

Chintalapati, S., Kiran, M. D., & Shivaji, S. (2004). Role of membrane lipid fatty acids in cold adaptation. *Cellular and molecular biology (Noisy-le-Grand*, France), 50(5), 631–642.

Chung, D. J., Sparagna, G. C., Chicco, A. J., & Schulte, P. M. (2018). Patterns of mitochondrial membrane remodeling parallel functional adaptations to thermal stress. The Journal of experimental biology, 221(Pt 7), jeb174458.

Chwastek, G., Surma, M. A., Rizk, S., Grosser, D., Lavrynenko, O., Rucińska, M., Jambor, H., & Sáenz, J. (2020). Principles of Membrane Adaptation Revealed through Environmentally Induced Bacterial Lipidome Remodeling. Cell reports, 32(12), 108165.

Clark, M. S., & Peck, L. S. (2009). Triggers of the HSP70 stress response: environmental responses and laboratory manipulation in an Antarctic marine invertebrate (Nacella concinna). Cell stress & chaperones, *14*(6), 649–660.

Clarke, A. (1980). The biochemical composition of krill, Euphausia superba Dana, from South Georgia. Journal of Experimental Marine Biology and Ecology, 43(3), 221–236.

Clarke, A. (1984). Lipid Content and Composition of Antarctic Krill, *Euphausia Superba* Dana, Journal of Crustacean Biology, 4(*5*), 285–294.

Crockett, E. L., & Sidell, B. D. (1993). Peroxisomal beta-oxidation is a significant pathway for catabolism of fatty acids in a marine teleost. The American journal of physiology, *264*(5 Pt 2), R1004–R1009.

Cuzin-Roudy, J., Irisson, J.-O., Penot, F., Kawaguchi, S., & Vallet, C. (2014). Southern Ocean Euphausiids. In C. De Broyer, P. Koubbi, H. J. Griffiths, B. Raymond, C. Udekem d’Acoz, A. P. Van de Putte, B. Danis, B. David, S. Grant, J. Gutt, C. Held, G. Hosie, F. Huettmann, A. Post, & M. Rauschert (Eds.), Biogeographic Atlas of the Southern Ocean (pp. 309-320). Scientific Committee on Antarctic Research.

Du, M., Chen, J., Wang, C., Jiang, Z., Wang, M., Pang, M., Bu, T., Cong, R., Wang, W., Zhang, G., & Li, L. (2025). Membrane Lipid Remodeling Strategies Regulate Fluidity for Acute Temperature Adaptation in Oysters. Evolutionary applications, 18(9), e70156.

Dunn, W. B., Broadhurst, D., Begley, P., Zelena, E., Francis-McIntyre, S., Anderson, N., Brown, M., Knowles, J. D., Halsall, A., Haselden, J. N., Nicholls, A. W., Wilson, I. D., Kell, D. B., Goodacre, R., Human Serum Metabolome (HUSERMET) Consortium (2011). Procedures for large-scale metabolic profiling of serum and plasma using gas chromatography and liquid chromatography coupled to mass spectrometry. Nature protocols, 6(7), 1060–1083.

Falabella, M., Vernon, H. J., Hanna, M. G., Claypool, S. M., Pitceathly, R. D. S. (2021). Cardiolipin, Mitochondria, and Neurological Disease. Trends in endocrinology and metabolism: TEM, 32(4), 224–237.

Feugere, L., Fernandes, J. F., Guscelli, E., Melo, T., Aveiro, S., Chabot, D., Calado, R., Domingues, R., Madeira, D., & Calosi, P. (2025). Non-additive phospholipidomic responses to ocean warming and acidification drive intraspecific variation in cell membrane vulnerability in a marine ectotherm. Environmental research, 283, 122135.

Flores H, Atkinson A, Kawaguchi S, Krafft BA, Milinevsky G, Nicol S, Reiss C, Tarling GA, Werner R, Bravo Rebolledo E, Cirelli V, Cuzin-Roudy J, Fielding S, Groeneveld JJ, Haraldsson M, Lombana A, Marschoff E, Meyer B, Pakhomov EA, Rombolá E, Schmidt K, Siegel V, Teschke M, Tonkes H, Toullec JY, Trathan PN, Tremblay N, Van de Putte, van Franeker JA, Werner T. (2012). Impact of climate change on Antarctic krill. Mar Ecol Prog Ser, 458:1–19.

Fraser, K. P. P., Peck, L. S., Clark, M. S., Clarke, A., & Hill, S. L. (2022). Life in the freezer: protein metabolism in Antarctic fish. Royal Society open science, 9(3), 211272.

Hagen, W., Kattner, G. (1998). Lipid metabolism of the Antarctic euphausiid *Thysanmssa* macrura and its ecological implications, Limnology and Oceanography, 43, 1894–1901.

Hagen, W., Kattner, G., Terbrüggen, A., Van Vleet, E. S. (2001). Lipid metabolism of the Antarctic krill *Euphausia superba* and its ecological implications. Marine Biology, 139, 95–104.

Hazel, J. R., & Williams, E. E. (1990). The role of alterations in membrane lipid composition in enabling physiological adaptation of organisms to their physical environment. Progress in lipid research, 29(3), 167–227.

He, A., Dean, J. M., & Lodhi, I. J. (2021). Peroxisomes as cellular adaptors to metabolic and environmental stress. Trends in cell biology, 31(8), 656–670.

Hellessey, N., Ericson, J. A., Nichols, P. D., Kawaguchi, S., Nicol, S., Hoem, N., Virtue, P. (2018). Seasonal and interannual variation in the lipid content and composition of *Euphausia superba* Dana, 1850 (Euphausiacea) samples derived from the Scotia Sea fishery. Journal of Crustacean Biology, *38(6)*, 673–681.

Heyen, S., Schneider, V., Hüppe, L., Meyer, B., & Wilkes, H. (2023). Variations of intact phospholipid compositions in the digestive system of Antarctic krill, *Euphausia superba*, between summer and autumn. PloS one, 18(12), e0295677.

Hiltunen, J. K., Kärki, T., Hassinen, I. E., Osmundsen, H. (1986). beta-Oxidation of polyunsaturated fatty acids by rat liver peroxisomes. A role for 2,4-dienoyl-coenzyme A reductase in peroxisomal beta-oxidation. The Journal of biological chemistry, *261*(35), 16484–16493.

Inigo, M., Deja, S., & Burgess, S. C. (2021). Ins and Outs of the TCA Cycle: The Central Role of Anaplerosis. Annual review of nutrition, 41, 19–47.

Jiao, S., Nie, M., Song, H., Xu, D., You, F. (2020). Physiological responses to cold and starvation stresses in the liver of yellow drum (Nibea albiflora) revealed by LC-MS metabolomics. The Science of the total environment, 715, 136940.

Ju, S. J., Harvey, H. R. (2004). Lipids as markers of nutritional condition and diet in the Antarctic krill Euphausia superba and Euphausia crystallorophias during austral winter, Deep Sea Research Part II: Topical Studies in Oceanography, 51(17–19), 2199–2214.

Ju, SJ., Kang, HK., Kim, W.S., Harvey, H. R. (2009). Comparative lipid dynamics of euphausiids from the Antarctic and Northeast Pacific Oceans. Marine Biology, 156, 1459–1473.

Kang, M. J., Fujino, T., Sasano, H., Minekura, H., Yabuki, N., Nagura, H., Iijima, H., & Yamamoto, T. T. (1997). A novel arachidonate-preferring acyl-CoA synthetase is present in steroidogenic cells of the rat adrenal, ovary, and testis. Proceedings of the National Academy of Sciences of the United States of America, 94(7), 2880–2884.

Kattner, G., & Hagen, W. (1998). Lipid metabolism of the Antarctic euphausiid *Euphausia crystallorophias* and its ecological implications. Marine Ecology Progress Series, 170, 203–213.

Kawaguchi, S., Atkinson, A., Bahlburg, D., Bernard, K. S., Cavan, E. L., Cox, M. J., Hill, S. L., Meyer, B., Veytia, D. (2024). Climate change impacts on Antarctic krill behaviour and population dynamics. Nature Reviews Earth & Environment, 5, 43–58.

Kuhl, C., Tautenhahn, R., Böttcher, C., Larson, T. R., Neumann, S. (2012). CAMERA: an integrated strategy for compound spectra extraction and annotation of liquid chromatography/mass spectrometry data sets. Analytical chemistry, 84(1), 283–289.

Leonardi, R., & Jackowski, S. (2007). Biosynthesis of Pantothenic Acid and Coenzyme A. EcoSal Plus, 2(2), 10.1128/ecosalplus.3.6.3.4.

Liu, S., Fu, S., Wang, G., Cao, Y., Li, L., Li, X., Yang, J., Li, N., Shan, Y., Cao, Y., Ma, Y., Dong, M., Liu, Q., & Jiang, H. (2021). Glycerol-3-phosphate biosynthesis regenerates cytosolic NAD^+^ to alleviate mitochondrial disease. Cell metabolism, 33(10), 1974–1987.e9.

Lubawy, J., Chowański, S., Adamski, Z., & Słocińska, M. (2022). Mitochondria as a target and central hub of energy division during cold stress in insects. Frontiers in zoology, 19(1), 1.

Lynes, M. D., Shamsi, F., Sustarsic, E. G., Leiria, L. O., Wang, C. H., Su, S. C., Huang, T. L., Gao, F., Narain, N. R., Chen, E. Y., Cypess, A. M., Schulz, T. J., Gerhart-Hines, Z., Kiebish, M. A., & Tseng, Y. H. (2018). Cold-Activated Lipid Dynamics in Adipose Tissue Highlights a Role for Cardiolipin in Thermogenic Metabolism. Cell reports, 24(3), 781– 790.

Madiraju, S. R. M., Possik, E., Al-Mulla, F., Nolan, C. J., & Prentki, M. (2026). Glycerol and Glycerol-3-Phosphate: Multifaceted Metabolites in Metabolism, Cancer, and Other Diseases. Endocrine reviews, 47(1), 93–120.

Mann H. B., Whitney D. R. (1947). On a Test of Whether one of Two Random Variables is Stochastically Larger than the Other. The Annals of Mathematical Statistics, 18(1), 50– 60.

Marszalek, J. R., Kitidis, C., Dirusso, C. C., & Lodish, H. F. (2005). Long-chain acyl-CoA synthetase 6 preferentially promotes DHA metabolism. The Journal of biological chemistry, 280(11), 10817–10826.

Mayzaud, P., Boutoute, M., & Alonzo, F. (2003). Lipid composition of the euphausiids *Euphausia vallentini* and *Thysanoessa macrura* during summer in the Southern Indian Ocean. Antarctic Science, 15(4), 463–475.

Meyer, B. (2012). The overwintering of Antarctic krill, *Euphausia superba*, from an ecophysiological perspective. Polar Biol, 35, 15–37.

Meyer, B., Auerswald, L., Siegel, V., Spahic, S., Pape, C., Fach, B. A., Teschke, M., Lopata, A. L., & Fuentes, V. (2010). Seasonal variation in body composition, metabolic activity, feeding, and growth of adult krill *Euphausia superba* in the Lazarev Sea. Marine Ecology Progress Series, 398, 1–18

Murphy, E. J., Watkins, J. L., Trathan, P. N., Reid, K., Meredith, M. P., Thorpe, S. E., Johnston, N. M., Clarke, A., Tarling, G. A., Collins, M. A., Forcada, J., Shreeve, R. S., Atkinson, A., Korb, R., Whitehouse, M. J., Ward, P., Rodhouse, P. G., Enderlein, P., Hirst, A. G., Martin, A. R., … Fleming, A. H. (2007). Spatial and temporal operation of the Scotia Sea ecosystem: a review of large-scale links in a krill centred food web. *Philosophical transactions of the Royal Society of London. Series B*, Biological sciences, 362(1477), 113–148.

Murzina, S. A., Voronin, V. P., Bitiutskii, D. G., Mishin, A. V., Khurtina, S. N., Frey, D. I., & Orlov, A. M. (2023). Comparative Analysis of the Fatty Acid Profiles of Antarctic Krill (*Euphausia superba* Dana, 1850) in the Atlantic Sector of the Southern Ocean: Certain Fatty Acids Reflect the Oceanographic and Trophic Conditions of the Habitat. Journal of Marine Science and Engineering, *11*(10), 1912.

Overgaard, J., Gerber, L., & Andersen, M. K. (2021). Osmoregulatory capacity at low temperature is critical for insect cold tolerance. Current opinion in insect science, 47, 38–45.

Pang, Z., Lu, Y., Zhou, G., Hui, F., Xu, L., Viau, C., Spigelman, A.F., MacDonald, P.E., Wishart, D.S., Li, S., & Xia, J. (2024). MetaboAnalyst 6.0: towards a unified platform for metabolomics data processing, analysis and interpretation. Nucleic Acids Research, 52, 398–406

Pauli, N. C., Flintrop, C. M., Konrad, C., Pakhomov, E. A., Swoboda, S., Koch, F., Wang, X. L., Zhang, J. C., Brierley, A. S., Bernasconi, M., Meyer, B., & Iversen, M. H. (2021). Krill and salp faecal pellets contribute equally to the carbon flux at the Antarctic Peninsula. Nature communications, 12(1), 7168.

Peck L. S. (2016). A Cold Limit to Adaptation in the Sea. Trends in ecology & evolution, 31(1), 13–26.

Phleger, C. F., Nelson, M. M., Mooney, B. D., & Nichols, P. D. (2002). Interannual and between species comparison of the lipids, fatty acids and sterols of Antarctic krill from the US AMLR Elephant Island survey area. Comparative biochemistry and physiology. Part B, Biochemistry & molecular biology, 131(4), 733–747.

Qu, L., Xiao, Y., Jia, Z., Wang, Z., Wang, C., Hu, T., Wu, C., Zhang, J. (2015). Comprehensive two-dimensional liquid chromatography coupled with quadrupole time-of-flight mass spectrometry for chemical constituents analysis of tripterygium glycosides tablets. Journal of chromatography. A, 1400, 65–73.

Ren, X., Yu, Z., Xu, Y., Zhang, Y., Mu, C., Liu, P., Li, J. (2020). Integrated transcriptomic and metabolomic responses in the hepatopancreas of kuruma shrimp (*Marsupenaeus japonicus*) under cold stress. Ecotoxicology and environmental safety, 206, 111360.

Ronges, D., Walsh, J. P., Sinclair, B. J., Stillman, J. H. (2012). Changes in extreme cold tolerance, membrane composition and cardiac transcriptome during the first day of thermal acclimation in the porcelain crab Petrolisthes cinctipes. The Journal of experimental biology, 215(Pt 11), 1824–1836.

Saether, O., Ellingsen, T. E., & Mohr, V. (1986). Lipids of North Atlantic krill. Journal of lipid research, 27(3), 274–285.

Saether, O., Ellingsen, T. E., Mohr, V. (1985). The distribution of lipid in the tissues of Antarctic krill, Euphausia superba. Comparative Biochemistry and Physiology Part B: Comparative Biochemistry, 81(*3*), 609–614.

Safronova, N., Junghans, L., & Saenz, J. P. (2024). Temperature change elicits lipidome adaptation in the simple organisms Mycoplasma mycoides and JCVI-syn3B. Cell reports, 43(7), 114435.

Savoca, M. S., Kumar, M., Sylvester, Z., Czapanskiy, M. F., Meyer, B., Goldbogen, J. A., & Brooks, C. M. (2024). Whale recovery and the emerging human-wildlife conflict over Antarctic krill. Nature communications, 15(1), 7708.

Sinensky M. (1974). Homeoviscous adaptation--a homeostatic process that regulates the viscosity of membrane lipids in Escherichia coli. Proceedings of the National Academy of Sciences of the United States of America, 71(2), 522–525.

Smith, A. J. R., Wotherspoon, S., Ratnarajah, L., Cutter, G. R., Macaulay, G. J., Hutton, B., King, R., Kawaguchi, S., & Cox, M. J. (2025). Antarctic krill vertical migrations modulate seasonal carbon export. *Science (New York*, N.Y.), 387(6732), eadq5564.

Smith, C. A., Want, E. J., O’Maille, G., Abagyan, R., Siuzdak, G. (2006). XCMS: processing mass spectrometry data for metabolite profiling using nonlinear peak alignment, matching, and identification. Analytical chemistry, 78(3), 779–787.

Stincone, A., Prigione, A., Cramer, T., Wamelink, M. M., Campbell, K., Cheung, E., Olin-Sandoval, V., Grüning, N. M., Krüger, A., Tauqeer Alam, M., Keller, M. A., Breitenbach, M., Brindle, K. M., Rabinowitz, J. D., & Ralser, M. (2015). The return of metabolism: biochemistry and physiology of the pentose phosphate pathway. Biological reviews of the Cambridge Philosophical Society, 90(3), 927–963.

Stübing, D., & Hagen, W. (2003). Fatty acid biomarker ratios—suitable trophic indicators in Antarctic euphausiids? Polar Biology, 26, 774–782.

Sustarsic, E. G., Ma, T., Lynes, M. D., Larsen, M., Karavaeva, I., Havelund, J. F., Nielsen, C. H., Jedrychowski, M. P., Moreno-Torres, M., Lundh, M., Plucinska, K., Jespersen, N. Z., Grevengoed, T. J., Kramar, B., Peics, J., Hansen, J. B., Shamsi, F., Forss, I., Neess, D., Keipert, S., … Gerhart-Hines, Z. (2018). Cardiolipin Synthesis in Brown and Beige Fat Mitochondria Is Essential for Systemic Energy Homeostasis. Cell metabolism, 28(1), 159–174.e11.

TeSlaa, T., Ralser, M., Fan, J., & Rabinowitz, J. D. (2023). The pentose phosphate pathway in health and disease. Nature metabolism, 5(8), 1275–1289.

Touchette, D., Altshuler, I., Gostinčar, C., Zalar, P., Raymond-Bouchard, I., Zajc, J., McKay, C. P., Gunde-Cimerman, N., Whyte, L. G. (2022). Novel Antarctic yeast adapts to cold by switching energy metabolism and increasing small RNA synthesis. The ISME journal, 16(1), 221–232.

Toullec, J. Y., Cascella, K., Ruault, S., Geffroy, A., Lorieux, D., Montagné, N., Ollivaux, C., & Lee, C. Y. (2020). Antarctic krill (Euphausia superba) in a warming ocean: thermotolerance and deciphering Hsp70 responses. Cell stress & chaperones, 25(3), 519–531.

Tovar-Sanchez, A., C. M. Duarte, S. Hernández-León, and S. A. Sañudo-Wilhelmy. (2007). Krill as a central node for iron cycling in the Southern Ocean, Geophysical Research Letters, 34, L11601.

Virtue, P., Nichols, P. D., Nicol, S., McMinn, A., Sikes, E. L. (1993b). The lipid composition of Euphausia superba Dana in relation to the nutritional value of Phaeocystis pouchetii (Hariot) Lagerheim. Antarctic Science, 5(02), 169–177.

Virtue, P., Nicol, S. & Nichols, P.D. (1993a). Changes in the digestive gland of *Euphausia superba* during short-term starvation: lipid class, fatty acid and sterol content and composition. Marine Biology, 117, 441–448.

Wang, J., Zhang, T., Shen, X., Liu, J., Zhao, D., Sun, Y., Wang, L., Liu, Y., Gong, X., Liu, Y., Zhu, Z. J., Xue, F. (2016). Serum metabolomics for early diagnosis of esophageal squamous cell carcinoma by UHPLC-QTOF/MS. Metabolomics, 12, 116.

Want, E. J., Wilson, I. D., Gika, H., Theodoridis, G., Plumb, R. S., Shockcor, J., Holmes, E., Nicholson, J. K. (2010). Global metabolic profiling procedures for urine using UPLC-MS. Nature protocols, 5(6), 1005–1018.

Wu, D., Huang, Y., Chen, Q., Jiang, Q., Li, Y., & Zhao, Y. (2019). Effects and transcriptional responses in the hepatopancreas of red claw crayfish *Cherax quadricarinatus* under cold stress. Journal of thermal biology, 85, 102404.

Wu, G., Baumeister, R., & Heimbucher, T. (2023). Molecular Mechanisms of Lipid-Based Metabolic Adaptation Strategies in Response to Cold. Cells, 12(10), 1353.

Yamashita, A., Hayashi, Y., Nemoto-Sasaki, Y., Ito, M., Oka, S., Tanikawa, T., Waku, K., & Sugiura, T. (2014). Acyltransferases and transacylases that determine the fatty acid composition of glycerolipids and the metabolism of bioactive lipid mediators in mammalian cells and model organisms. Progress in lipid research, 53, 18–81.

